# Fiber-associated *Lachnospiraceae* reduce colon tumorigenesis by modulation of the tumor-immune microenvironment

**DOI:** 10.1101/2021.02.24.432654

**Authors:** Ana S Almeida, Tam T T Tran, Tarini S. Ghosh, Celine Ribiere, Cathriona Foley, Lisa A Azevedo, Paola Pellanda, Werner Frei, Cara M Hueston, Raju Kumar, Burkhardt Flemer, Inês Sequeira, Micheal O’Riordain, Fergus Shanahan, Paul W. O’Toole

## Abstract

Patients with colorectal cancer (CRC) harbor gut microbiomes that differ in structure and function from those of healthy individuals, suggesting this altered microbiome could contribute to tumorigenesis. Despite increasing evidence implicating the gut microbiome in CRC, the collective role of different microbial consortia in CRC carcinogenesis is unclear. We have previously described these consortia as co-abundance groups that co-exist at different abundance levels in the same patient. Here, we report that tumor biopsy tissue from patients with a “high-risk” Pathogen-type microbiome had a different immune transcriptome and immune cell infiltrate from those with a “low-risk” *Lachnospiraceae*-type microbiome. Transplantation from patients of the two fecal microbiome types into mice with an orthotopic tumor differentially affected tumor growth and the systemic anti-tumor immune response. The differences in tumor volume and immunophenotype between mice receiving the high-risk and the low-risk microbiome correlated with differences in the engrafted human microbial species and predicted microbiome-encoded metabolites in the two groups. Of twelve taxa whose abundance in recipient mice led to increased tumor onset, seven corresponded with differentially abundant taxa in a global dataset of 325 CRC patients versus 310 healthy controls. These data suggest that the enrichment for a *Lachnospiraceae*-type configuration of the gut microbiome may influence colon cancer progression and disease outcome by modulating the local and systemic anti-tumor immune response.

**Graphical Abstract:** 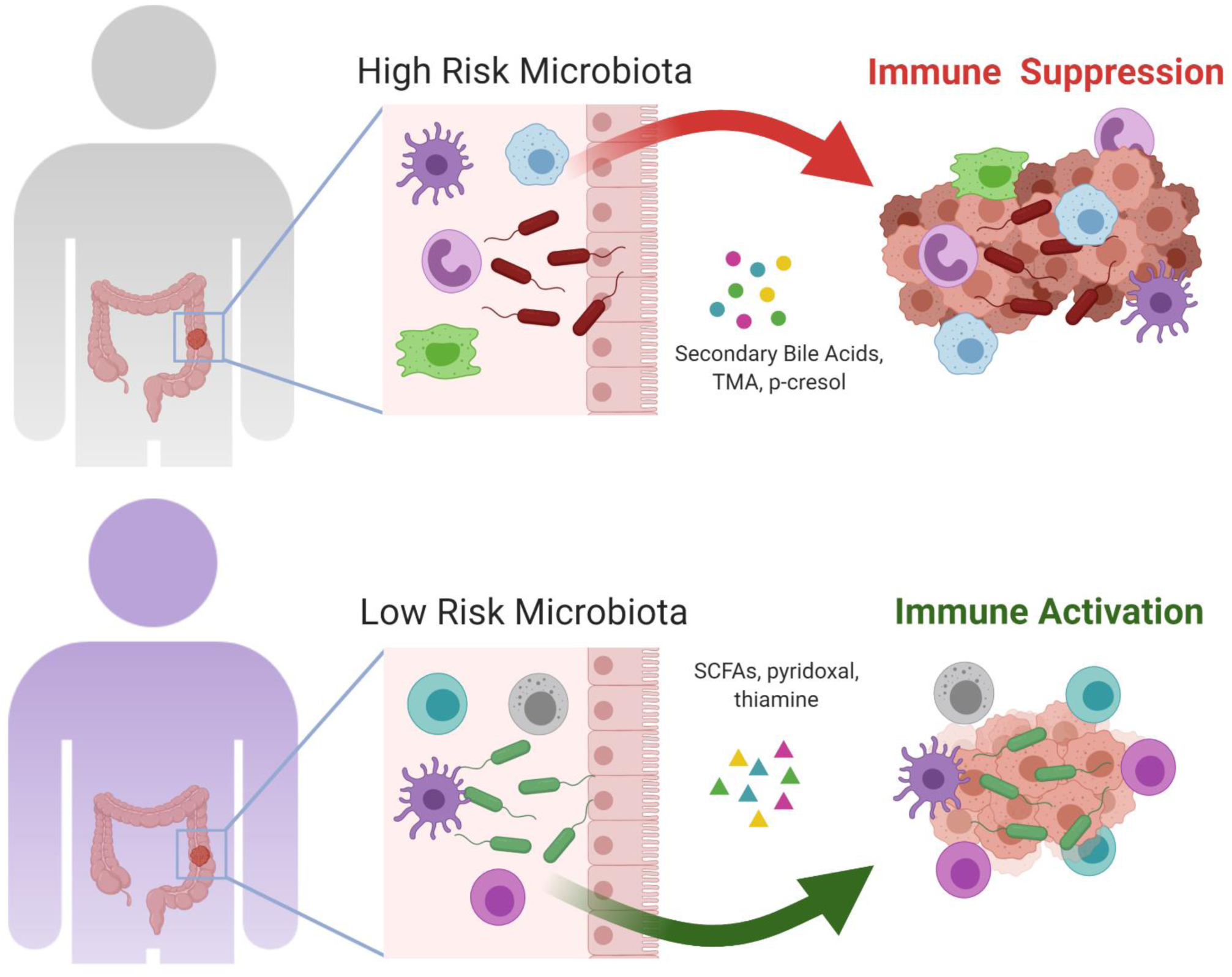

Proposed model of how the high-risk Pathogen and low-risk *Lachnospiraceae* CAGs differentially modulate the tumor immune response.

## Introduction

Colorectal cancer (CRC) is among the top three causes of global cancer-related mortality and the incidence continues to increase worlwide^1^. Risk factors for CRC include age, a diet low in fiber and rich in red meat and fat, and chronic inflammation of the gastrointestinal tract^2, 3^. All these factors are closely associated with altered composition and function of the gut microbiota. The gut microbiota from patients with polyps or CRC differs from that of healthy individuals^4, 5^. However, whether this “dysbiosis” contributes to disease or is a consequence of cancer is not yet clear^6^. Differences in the gut microbiota also explain why some patients with cancer benefit from new cancer immunotherapies while other patients do not^7^.

Studies to date in mice have indicated several ways by which specific bacterial species might impact CRC development. Proposed CRC-promoting bacteria or “oncobacteria” include *Fusobacterium nucleatum*^8, 9^, genotoxic (*pks*+) *Escherichia coli*^10, 11^, enterotoxigenic *Bacteroides fragilis*^12^, *Streptococcus gallolyticus*^13^, and *Peptostreptococcus anaerobius*^14^. Despite the increasing evidence implicating the gut microbiota in CRC development, the role of microbial community-driven pathogenicity as distinct from single taxa still needs to be elucidated.

Bacteria can directly contribute to the development of CRC by the release of genotoxic factors^15^, production of carcinogenic metabolites^16^, or by interacting with the immune system^17^. Infiltration by immune cells heavily impacts clinical outcome in CRC^18^. Numbers of intratumoral T cells correlate positively with better CRC outcomes, including disease-free and overall survival^19^. However, the role of innate immune cells in CRC is less clear. Under certain circumstances, neutrophils and macrophages release radical oxygen species and nitrous oxide, which can potentially cause genomic damage in colonic epithelial cells^20^. The presence of specific immune cells in the tumor microenvironment, as well as the release of lymphocyte-attracting chemokines and cytokines, can be modulated by the microbiota, either directly or through their metabolites^6^. Understanding how specific microbiota taxa and their metabolites differentially modulate the host immune response has important clinical implications for CRC patients, including diagnosis, and potentially also treatment.

In our previous analyses of the gut microbiome in CRC patients, we used clustering methodologies to identify six co-abundance groups or CAGs (i.e. co-abundance associations of genera)^4^, which we subsequently simplified to five CAGs^21^. A single subject harbored multiple CAGs but their relative abundance differed between CRC patients and healthy controls, with three of the 5 CAGs being more abundant in CRC patients. Tumor biopsies from patients whose microbiome was dominated by these CAGs showed differential expression of 18 genes involved in inflammation and CRC progression^4^, suggesting a possible microbiome influence on tumor development. To test this, here we used human-into-mice fecal microbiota transplants from patients with adenomas or adenocarcinomas with a ‘high risk’ or a ‘low risk’ microbiota. We show for the first time that the presence of a Pathogen CAG or a *Lachnospiraceae* CAG microbiome differentially affected tumor progression. Distinct bacterial taxa correlated with tumorigenesis, different metabolic pathways, and divergent systemic immune responses that correlated with tumor volume.

## Results

### Different global transcriptome in human colonic tumors with the Pathogen CAG and the *Lachnospiraceae* CAG

To investigate if bacterial CAGs present in CRC patients may shape the tumor immune profile and affect the host response, we recruited a cohort of 32 treatment-naive patients with either adenomas or adenocarcinomas and undergoing resection surgery **(Fig. 1A, Table S1**). None of these patients had received antibiotics or probiotics in the 3 months before surgery, nor been treated with chemotherapy or radiation therapy. Samples were collected from different mucosal sites, including from the tumor (or adenoma) and nearby healthy tissue. Using the approach we previously described^4, 22^, we profiled the abundance of the 5 different bacterial CAGs in the 32 patients and named them according to the dominant genera in each group –-*Bacteroidetes*, *Lachnospiraceae*, Pathogen, *Prevotella*, and *Rumminococcus* (**Figs. 1B, S1, S2; Table S2**). As expected, the majority of the patients had a higher abundance of the Pathogen CAG compared to the other CAGs.

**Figure 1.**
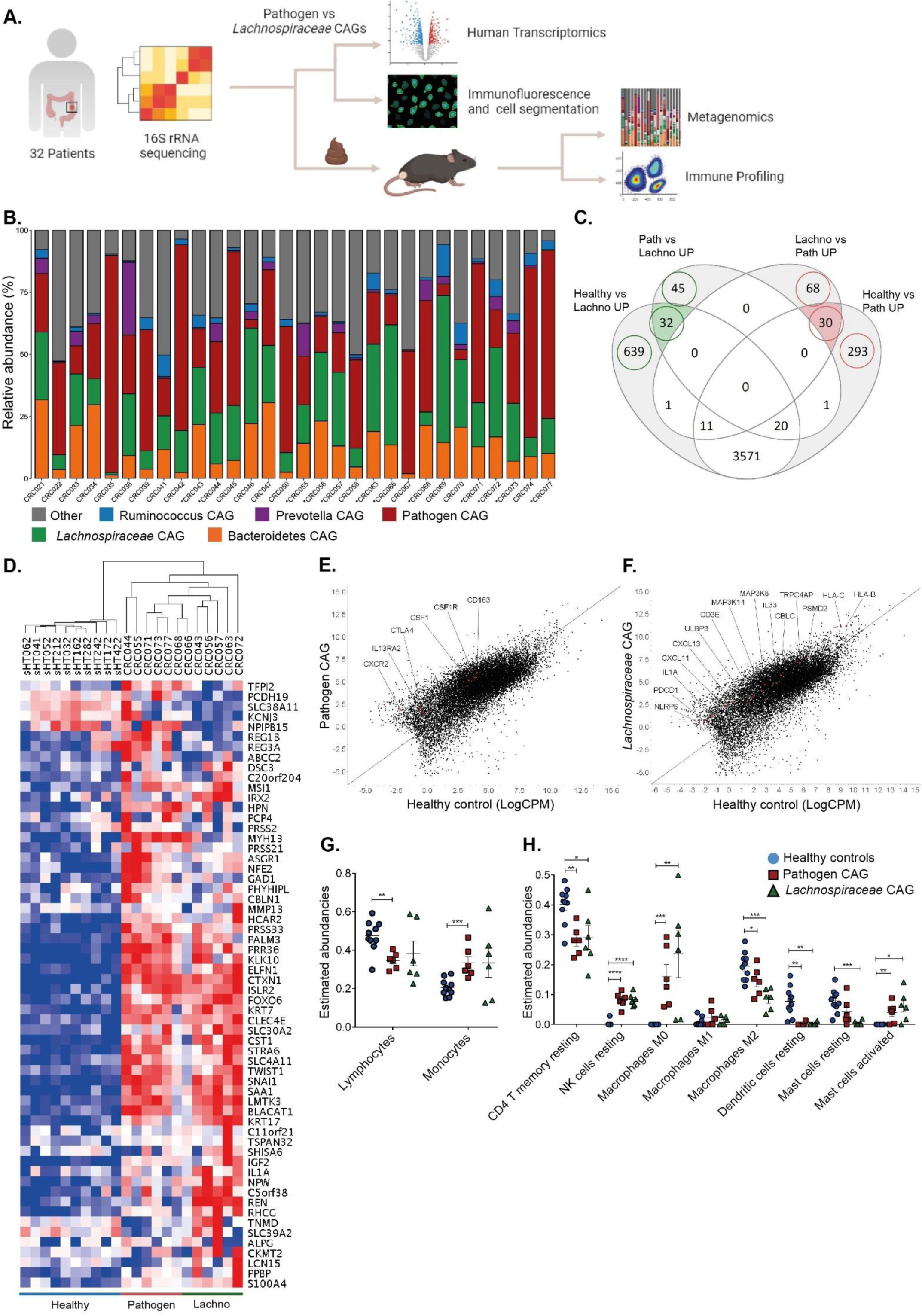
Distinct microbiomes in patients with adenomas and CRC correlate with differential human immune transcripts. **A.** Overview of the experimental design. 32 treatment-naïve patients were included in the study. Surgical resections were collected from multiple sites in the colon and analyzed by 16S rRNA sequencing, immunofluorescence, and RNA expression analysis. Selected fecal samples were collected anaerobically and administered in a germ-free cancer mouse model. **B.** Human microbiota composition measured by proportional abundance of bacterial CAGs in human colon biopsies. CAGs are named after the most abundant genus. Stars (*) indicate the 12 patients selected for bulk RNAseq analysis. **C.** Venn diagram depicting numbers of significantly DEGs (*p* value <0.5) between healthy controls (Healthy, n= 10), tumors from the Pathogen CAG (Path, n= 6), and tumors from subjects harboring the *Lachnospiraceae* CAG (Lachno, n= 6). The gene numbers circled in red are those uniquely elevated in the Pathogen CAG and gene numbers circled in green are uniquely elevated in the *Lachnospiraceae* CAG. **D.** Heatmap of unsupervised hierarchal clustering of genes and patients, representing the top 60 significantly DEGs (CRC versus healthy controls; FDR adjusted *p* value < 0.1, and *Lachnospiraceae* CAG versus Pathogen CAG; *p* value < 0.05. All log_2_FC ≤-1.5 and ≥1.5), consisting of the top 10 significantly DEGs from each circle highlighted in the Venn diagram (panel C). **E, F.** Expression plots displaying labeled immune genes differentially expressed and uniquely elevated in (E) Pathogen CAG and (F) *Lachnospiraceae* CAG tumors relative to healthy controls. The x-axis is the logCPM values for healthy controls and the y-axis is the logCPM values for (E) Pathogen CAG and (F) *Lachnospiraceae* CAG tumors. Red dots show genes of interest. **G, H**. Estimated immune cell abundancies from whole transcriptomic data deconvoluted with the CIBERSORTx software in healthy controls and tumors. Estimated lymphocyte abundances were calculated as the sum of proportions of naïve B cells, memory B cells, CD8^+^ T cells, naïve CD4^+^ T cells, resting memory CD4^+^ T cells, and activated memory CD4^+^ T cells. Estimated monocyte abundances were calculated as the sum of proportions of monocytes, M0 macrophages, M1 macrophages, and M2 macrophages. Bars represent mean ±SEM. *p* values were calculated by unpaired Student’s t-tests. \**p* ≤ 0.05, \*\**p* ≤ 0.01, \*\*\**p* ≤ 0.001, \*\*\*\**p* ≤ 0.0001.

**Figure 2.**
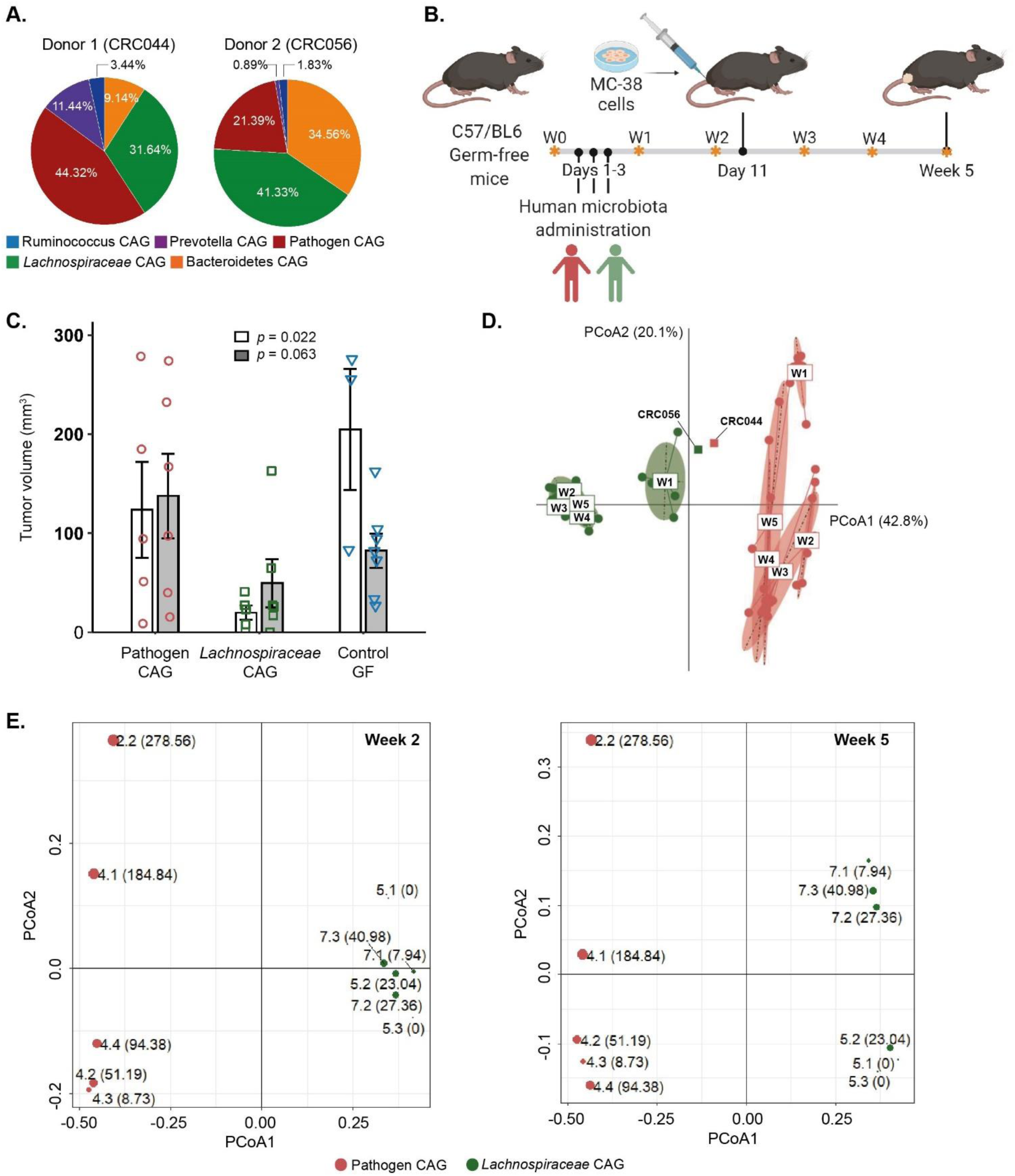
Tumor growth is strongly dependent on microbiota type in a humanized mouse model of CRC. **A.** Validation of microbiota composition in human donors for the pre-clinical mouse trial. Colon resections were collected at surgery and the mucosal microbiota was profiled using 16S rRNA sequencing. Pie-charts represent the abundance of the five bacterial CAGs on adenoma or tumor samples from each donor. Donor 1 (CRC044), diagnosed with a T3N2 rectum adenocarcinoma, was selected based on the high abundance of a Pathogen CAG microbiota and Donor 2 (CRC056), diagnosed with a tubulovillous adenoma, was selected based on the high abundance of a *Lachnospiraceae* CAG microbiota. **B.** Experimental design of the pre-clinical trial with a humanized MC-38 model of CRC. **C.** Tumor growth is reduced in mice receiving the *Lachnospiraceae* microbiota compared to mice receiving the Pathogen CAG microbiota or control germ-free (GF) mice. Tumor volume was measured with a caliper at endpoint and calculated as (length x width^2^)/2. Overall *p* values were calculated with the Kruskal-Wallis test. Data indicate mean ±SEM. n= 3-6 replicates/group per condition. Data from two independent experiments are shown by open and grey bars. **D.** Relatedness (β-diversity) of the fecal microbiota of the two human donors and respective recipient mice at different time-points represented by principal coordinate analysis (PCoA) on Bray-Curtis distance matrix (PERMANOVA r^2^=0.79; *p* value = 0.001). W, week. **E** PCoA plots of the Metaphlan2 species-level taxa profiles of the murine fecal microbiomes, performed at week 2 (left) and week 5 (right). Fecal microbiome profiles corresponding to the Pathogen CAG donor and the *Lachnospiraceae* CAG donor are colored in red and green, respectively. Each point corresponds to a specific mouse ID and the corresponding tumor volume is shown within parentheses. The size of each point is proportional to the tumor volume. PERMANOVA r^2^=0.24; *p* = 0.016 (week 2), r^2^=0.25; *p* value = 0.006 (week 5).

We selected 12 representative patients whose microbiome was dominated by either the Pathogen CAG (n= 6) or the *Lachnospiraceae* CAG (n= 6), together with biopsies from 10 healthy controls^4^, and subjected them to RNA seq analysis. Patients with adenomas or CRC were chosen accordingly to their microbiota profile and independently of tumor staging, size or location, although the majority of the tumors were located in the rectum or sigmoid colon (**Table S1**). First, we identified 21 (FDR <0.1) and an additional 518 (*p* value <0.05) genes significantly differentially expressed between Pathogen and *Lachnospiraceae* enriched tumors, with 268 upregulated in the Pathogen CAG and 271 upregulated in the *Lachnospiraceae* CAG tumors (**Fig. 1C, Tables S3 and S4**). Next, we identified significantly differentially expressed genes (DEGs) between healthy controls and the Pathogen CAG (11,520), and healthy controls and the *Lachnospiraceae* CAG (11,205) enriched tumors (**Fig. 1C, Table S3**). To further refine the transcriptomic differences found, we identified genes uniquely elevated in Pathogen or *Lachnospiraceae* dominated tumors compared to healthy controls and performed unsupervised hierarchical clustering (**Fig 1D**). This revealed that the two microbiota CAGs were associated with distinct host transcriptome differences. Pathway level analysis indicated that the greatest transcriptional changes were related to upregulation of the pathways involved in epithelial-mesenchymal transition, inflammatory response, angiogenesis, and interleukin-6 (IL-6) JAK STAT3 signaling in Pathogen-enriched tumors versus *Lachnospiraceae*-enriched tumors (**Fig. S3, Table S4**). We next focused on genes associated with the host immune response. Relative to healthy controls, *Lachnospiraceae* enriched tumors from patients with adenomas or CRC showed upregulated genes associated with immune signatures that were not different in Pathogen enriched tumors. These included antigen presentation genes (*HLA-C, HLA-B, TRPC4AP, PSMD2, ULBP3*), TCR signaling genes (*MAP3K8, CBLC, CD3E, MAP3K14*), chemokines predictive of increased survival in CRC^23, 24^ (*CXCL11, CXCL13*), innate immune genes (*IL1A, IL33, NLRP6*) and the inhibitory receptor PD1 (*PDCD1*) (**Fig. 1F**). In contrast, genes associated with monocytes or monocytic myeloid-derived suppressor cells (MDSCs) signature and signaling were differentially upregulated only in Pathogen-enriched tumors compared to healthy controls (CSF1R, CD163, CSF1, CXCR2), as well as the inhibitory receptor CTLA4 (**Fig. 1E**). These differentially expressed immune genes were also significant after testing for confounding by tumor site, tumor stage, and lymph node involvement (**Table S5**).

To functionally interpret these results, we used the CIBERSORTx (<u>C</u>ell type Identification <u>B</u>y <u>E</u>stimating <u>R</u>elative <u>S</u>ubsets <u>O</u>f known <u>R</u>NA <u>T</u>ranscripts) method^25, 26^ to deconvolute the gene expression data and estimate immune cell composition in CRC tumors. Pathogen CAG enriched tumors had a significantly higher estimated abundance of monocytes and lower estimated abundance of lymphocytes compared to healthy controls, whereas there were no statistical differences between *Lachnospiraceae* CAG tumors and healthy controls (**Fig 1G, Fig. S4, Table S4**). Specific leukocyte subset analysis identified a significant increase of M2 macrophages in Pathogen enriched tumors compared to *Lachnospiraceae* enriched tumors, but not M0 or M1 macrophages (**Fig. 1H**). Altogether these results suggest that the abundance of different microbiota taxa is associated with specific host gene expression changes, including differential upregulation of several immune pathways and recruitment of specific immune cells to the tumor microenvironment.

### A *Lachnospiraceae*-dominant microbiome reduced tumor growth in a pre-clinical mouse model of CRC

Although previous studies have determined that certain bacterial species can influence tumor progression and response to therapy^6, 7^, the mechanism(s) by which specific bacterial clusters present in CRC patients might influence tumor progression has not yet been determined. We therefore investigated the effect of the two different CAGs – Pathogen and *Lachnospiraceae* - on tumor development using the MC-38 orthotopic cancer mouse model^27^. This orthotopic model allows for the reliable and efficient study of tumors arising in immune-competent animals at the appropriate primary site^28^. Germ-free (GF) mice were administered one of two representative human microbiota types, from donors selected based on their mucosal microbiota profile (**Fig. 2A**). We selected donor 1 (CRC044, female, T3N2b rectum adenocarcinoma) because their mucosal microbiota was mostly composed of the Pathogen and *Prevotella* CAGs (44.3% and 11.4%, respectively, **Figs. 2A, S5, Table 1**). We selected donor 2 (CRC056, male, adenoma with low-grade dysplasia) with a mucosal microbiota dominated by the *Lacnhospiraceae* CAG (41.3%, **Figs. 2A, S5, Table 1**). After human fecal administration, we allowed 8-10 days for colonization and stabilization of the human microbiota^29^ and then injected the MC-38 colorectal cancer cells in the rectal submucosa of these mice, as previously described^30^ (**Fig. 2B**). Mice that received the Pathogen CAG microbiota developed larger tumors than mice that received the *Lachnospiraceae* CAG microbiota (**Fig. 2C**). Surprisingly, mice that received the *Lachnospiraceae* CAG developed very small tumors, even smaller when compared to the GF control group, and 3 out of 12 mice did not develop any tumors (*p*= 0.022 and *p*= 0.063, two independent experiments) (**Fig. 2C**). We also observed that mice that received the Pathogen CAG displayed a much higher level of variability in their tumor volume than the *Lachnospiraceae* CAG-receiving group.

**Table 1.** Clinico-pathological characteristics of the two donors.

|  | **CRC044** | **CRC056** |
| --- | --- | --- |
| **Gender** | Female | Male |
| **Age (years, at the time of surgery)** | 66 | 65 |
| **Tumor Type** | Adenocarcinoma | Tubulovillous adenoma with low grade dysplasia |
| **Tumor Location** | Rectum | Hepatic flexure |
| **Tumor Size (cm)** | 3.5 | - |
| **T-category** | T3 | T0 |
| **N-category** | N2b | - |
| **BMI** | N/A | 25.7 |
| **Hip/waist ratio** | N/A | 0.98 |
| **Medication** | Bendroflumethiazole 5mgs OD | Atrovasin 20mg OD, Eltroxin 50mgs OD |
| **Bowel Preparation** | Klean Prep x 1 and Picolax (5 days prior to surgery) followed by low residue diet x 5days and Phosphate enema x 2 on morning of the surgery | Klean Prep x 1 and Picolax (5 days prior to surgery) followed by low residue diet x 5days and Phosphate enema x 2 on morning of the surgery |
| **3-year clinical follow-up** | Patient had adjuvant chemotherapy following surgery. Patient finished oral Capecitabine in October 2017 and a CT scan in January 2018 which showed no evidence of disease. Development of symptoms (dysphasia and right-sided facial droop) in February 2020. 18 mm brain metastases consistent with rectal cancer. Negative CT scan and negative colonoscopy in June 2020. | No further treatment required to date |

We investigated the recipient mouse fecal microbiota composition using 16S rRNA gene amplicon (16S) sequencing. As expected, β-diversity analysis showed a donor-specific signature in each recipient group (R^2^= 0.79, *p*= 0.001) (**Fig. 2D**). Similar to β-diversity, α-diversity analysis of the recipient mice resembled donor microbiota diversity, with mice in the Pathogen CAG group harboring a less diverse fecal microbiota than the *Lachospiraceae* CAG group (**Fig. S6, S7, Table S6**). Strikingly, the differences observed in tumor growth between the Pathogen CAG and the *Lachnospiraceae* CAG mouse groups were significantly correlated with the mouse microbiota profiles at both weeks 2 (after human microbiota administration but before MC-38 cells injection) and week 5 (three weeks after MC-38 cells injection, endpoint) (R^2^= 0.25, *p*= 0.006) (**Figs. 2E**). This finding strengthened the causal link between the engrafted microbiota composition and tumor onset in the MC-38 mouse model. Notably, mice that received the Pathogen CAG were also characterized by significantly higher microbiome composition variability from the corresponding donor compared to those that received the *Lachnospiraceae* CAG, suggesting differences in the resilience or community cohesion of the two different microbiome types (**Fig. S8**).

### Distinct pathobiont and commensal taxa strongly correlate with tumorigenesis and predict tumor growth

To identify with greater taxonomic resolution the bacterial taxa that were associated with tumorigenesis, and their respective functions, we performed shallow shotgun sequencing^31^ on mouse fecal samples collected at weeks 2 and 5. Using Spearman’s correlation analysis between the abundance of the bacterial species present at each time point and the tumor volume at week 5, we identified several bacterial species that significantly correlated with tumor volume (**Figs. 3, S9**). Bacterial species that were positively correlated with tumor growth (referred to henceforth as ‘tumor positive’) were consistently present in higher abundance in mice that received the Pathogen CAG compared to mice that received the *Lachnospiraceae* CAG, while species negatively correlated with tumor growth (‘tumor negative’), showed the opposite trend. The taxa identified by 16S sequencing as tumor-positive or tumor-negative largely overlapped between the two experiments performed (Fig. S10), but the shallow shotgun sequencing data performed for the second replicate (experiment 2) afforded much greater taxonomic resolution and correspondingly more taxa. At week 2, we identified 23 taxa that showed significant association with tumor volume (**Fig. 3**). Interestingly, 8 out of these 23 were also associated with tumor volume at week 5 (**Fig. 3**). These included four tumor-negative bacterial taxa, 2 of which belonged to the *Lachnospiraceae* family (*Coprococcus comes* and *Ruminococcus lactaris*), and 4 tumor-positive taxa (*Ruminococcus obeum*, *Clostridium hathewayi, Flavonifractor plautii*, and *Coprococcus sp ART55 1*). In addition to these eight, there were 28 additional bacterial taxa whose abundance was significantly associated with tumor volume at week 5 only (**Fig. 3**). Tumor-positive taxa included *Bacteroides sp* (*B. fragilis*, *B. salyersiae, B. faecis, B. uniformis*), *Paraprevotella* sp., *Clostridium boltae,* and *Desulfovibrio piger*. Bacterial species that were negatively associated with tumor growth at week 5 included *Akkermansia muciniphila*, *Barnesiella intestinihominis*, *Alistipes* sp., and *Bifidobacterium* sp., (*B. longum* and *B. pseudocatenulatum*) (**Fig. 3**).

**Figure 3.**
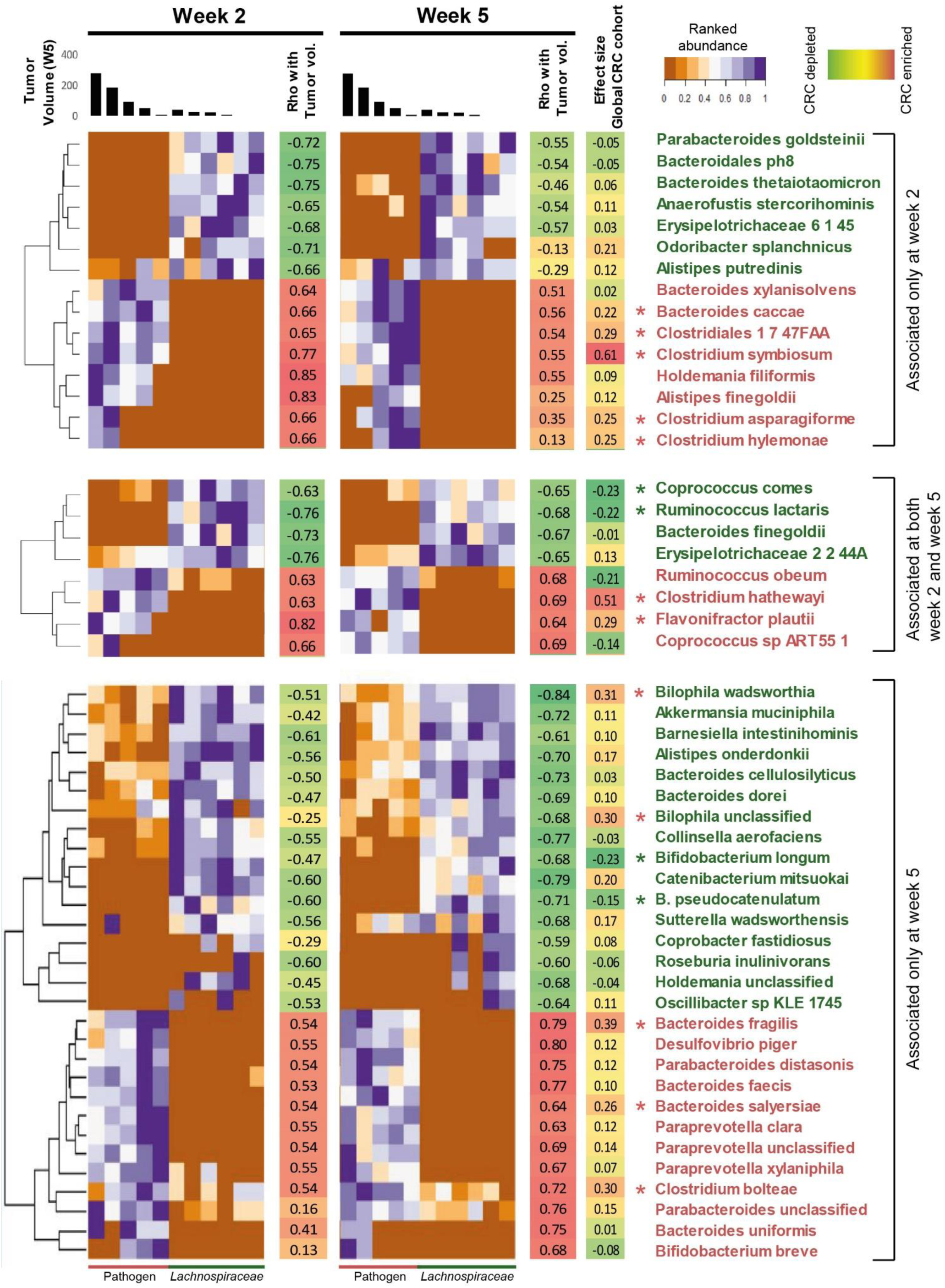
Distinct bacterial taxa are associated with final tumor volume. Heatmap showing the ranked abundances of ‘tumor positive’ (red) and ‘tumor negative’ (green) taxa in mice fecal microbiomes at week 2 only (top panel), shared between week 2 and 5 (middle panel), and week 5 only (bottom panel), as determined by shallow shotgun sequencing of samples from experiment 2. For each mouse, the corresponding tumor volume at week 5 is indicated as bar plots at the top of the heatmap. Spearman’s rank correlations between bacterial taxa abundance at different time points and the tumor volume at week 5 are indicated. The effect sizes (Cohen’s D) observed for the various taxa in the Global Reference cohort are also shown. Taxa that are significantly enriched or depleted in CRC (n = 325) versus healthy individuals (n = 310) (identified using Mann-Whitney U tests, *p* values corrected using Benjamini-Hochberg for FDR < 0.1) are indicated by red and green stars, respectively. Positive association with tumor volume and enrichment in CRC in the Global Reference cohort is color-coded in red. Negative association with tumor volume and depletion in CRC in the Global Reference cohort is color-coded in green.

To check the clinical relevance of these microbiome-tumor associations, we examined the abundance of these taxa in six patient-control paired studies whose CRC-associated microbiome had been shotgun sequenced (Global Reference CRC cohort, as described in the methods)^32–37^. We calculated the correlation between these bacterial taxa and CRC by comparing the effect size between CRC patients (n= 352) and healthy individuals (n= 310) across the six studies (**Fig. 3, Table S7**). We found 7 out of 12 tumor-positive taxa at week 2 to be significantly more abundant in CRC patients when compared to healthy subjects (**Fig. 3**). These included several Clostridia reported as pathobionts (*C. hylemonae*, *C. symbiosum*, *C. asparagiforme*, *C. hathewayi,* and *Clostridiales bacterium 1-7-47FAA*), along with *F. plautii* and *Bacteroides caccae*. In contrast, while 5 of the tumor-negative taxa were enriched in the healthy individuals, only *C. comes* and *R. lactaris* (*Lachnospiraceae* family) reached significance. Interestingly, *B. longum* and *B. pseudocatenulatum* (week 5) were also present in significantly lower abundance in CRC patients compared to healthy individuals, while *B. fragilis*, *B. salyersiae,* and *C. boltae* were significantly more abundant in CRC patients Some associations between bacterial taxa and tumor volume (e.g. *Bilophila* wadsworthia and *Bilophila* unclassified) were not confirmed in the human dataset. This suggests that these bacterial species might be donor-specific and not involved in tumor progression, at least in the MC-38 mouse model. Another example was *R. obeum* which showed a positive association with tumor volume at both weeks 2 and 5, but a negative association in the Global Reference CRC cohort. However, within the Pathogen CAG group, *R. obeum* showed a negative association with tumor volume, being in much higher abundance in mice that developed the smallest tumors (**Fig. S11**).

To explore if the dynamic shifts in microbial composition observed before tumor development (week 2) and afterward (week 5) could be a predictor of disease, we used Random-Forest (RF) classifier models to identify the bacterial species at the different time-points that predicted tumor volume. We found that the bacterial species at week 2 had a significantly higher predictive value compared to the bacterial species found at week 5 (*p* < 0.006) (**Figs. 4A, S12**). We also confirmed this effect was not due to a loss of species over time (data not shown). This result suggests that the transplanted taxa that correlated with tumor volume after 2 weeks of engraftment were better indicators of final tumor volume than the 5-week taxa, indicating that early microbe-cell interactions were pivotal for determining cancer progression.

**Figure 4.**
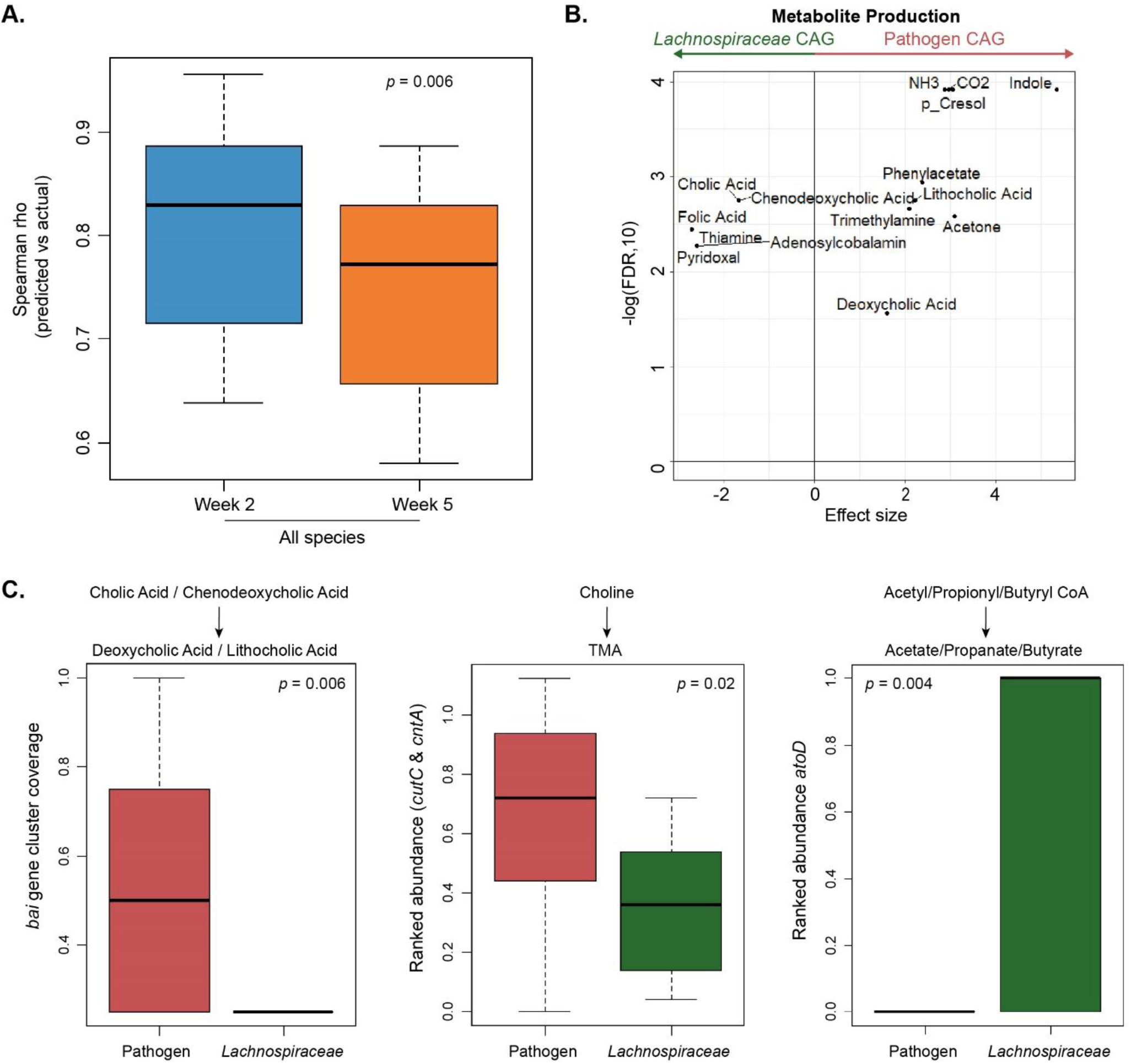
High-risk and low-risk microbiome are associated with different metabolic pathways. **A.** Pre-tumor microbiota is predictive of tumor growth. Tumor-associated bacterial taxa at week 2 have higher predictability for tumor volume than taxa at week 5. Boxplots show the variation of Spearman rho values calculated between the predicted and actual tumor volumes obtained for the 100 iterations of the two variants of RF models (trained on week 2 and week 5 abundance profiles, respectively). Mann-Whitney U test *p* values for the different comparisons are indicated. **B.** Volcano plot showing the validated set (identified as summarized in Fig. S13) of metabolite production functionalities that were predicted to have either a significant positive or negative association with the Pathogen CAG microbiome. The x-axis indicates the effect size difference (negative indicating enriched in the *Lachnospiraceae* CAG and positive indicating enriched in the Pathogen CAG), and the y-axis indicates the negative log of FDR value. **C.** Boxplots comparing (left) the coverage of the bile acid inducible (*bai*) gene cluster that converts the primary bile acids (cholic acid and chenodeoxycholic acid) into secondary bile acids (deoxycholic acid and lithocholic acid); (middle) the cumulated gene abundances of CntA and CutA enzymes that catalyze trimethylamine (TMA) production; and, (right) the abundance of the AtoD enzyme catalyzing the last step of short-chain fatty acids formation, between the Pathogen and the *Lachnospiraceae* microbiome types. The *p* values obtained using the Mann-Whitney U tests are indicated.

### The tumorigenic high-risk metagenome is characterized by detrimental metabolic pathways associated with CRC pathogenesis

The results obtained up to this point in our investigation strongly suggested a causal link between the microbiome type and tumorigenesis. To explore potential mechanisms, we investigated the metabolic functions associated with the two microbiota types. For this purpose, we identified a validated set of predicted metabolic capabilities that were 1) differentially abundant between the Pathogen CAG and *Lachnospiraceae* CAG mouse groups, and 2) differentially abundant in CRC patients versus healthy individuals in the Global Reference CRC cohorts (refer to **Fig. S13** for methodology). As expected, Pathogen CAG and *Lachnospiraceae* CAG microbiomes were associated with different metabolic pathways (Fig. S14). Differences in the predicted microbiome production of secondary bile acids (i.e. lithocholic acid (LCA) and deoxycholic acid (DCA)), trimethylamine (TMA), p-cresol, acetone, and ammonia were positively associated with the Pathogen CAG (**Fig. 4B**). Similarly, increased consumption of multiple short-chain fatty-acids (SCFAs), mainly driven by *D. piger*, was associated with the high-risk Pathogen CAG, a trait that could lead to the depletion of available SCFAs (**Fig. S15, Table S7**). In contrast, the predicted production of several vitamins including pyridoxal phosphate, folate, and thiamine, was associated with the low-risk *Lachnospiraceae* microbiota, exclusively driven by *B. longum* and *B. pseudocatenulatum* (**Fig. 4B, Table S8**). Interestingly, pyridoxal, an active form of vitamin B6, has been associated with a 30-50% reduction in CRC incidence^38^.

A limitation of the above data was that these were inferred based on experimentally known production and consumption profiles derived from reference genomes and organisms. To investigate whether these metabolic functionalities were also encoded in the genome of the strains present in the mouse fecal samples, we profiled the genes for enzymes known to confer these functions, focusing on the production of secondary bile acids, and genes for choline to TMA production. For secondary bile acid synthesis (DCA and LCA), a group of the *bai* gene cluster (*baiF*, *baiN*, *baiE*, *baiCD,* and *baiA)* is known to be involved. Based on the gene family abundances obtained in the HUMAnN2 analysis, we first checked and compared the coverage of the *bai* gene cluster in the samples derived from the Pathogen and the *Lachnospiraceae* microbiome recipient mice (**Fig. 4C**). The high-risk Pathogen CAG had significantly greater coverage of the *bai* gene cluster compared to the *Lachnospiraceae* CAG indicating a significantly greater probability of this metabolic function being present in the former (*p* < 0.006). Similarly, the Pathogen CAG also had a significantly larger copy number of the CutC and CntA enzymes that catalyze the conversion of choline to TMA, further validating the results of the inferred metabolite profiling (*p* < 0.02; **Fig. 4C**). In contrast, the low-risk *Lachnospiraceae* CAG had significantly higher copy numbers of the AtoD enzyme that catalyzes the formation of the anti-inflammatory SCFAs - butyrate, propionate, and acetate - from Butyryl CoA, Propionyl CoA, and Acetyl CoA (*p* < 0.0004 **Fig. 4C**).

### A *Lachnospiraceae*-type microbiome is associated with a strong systemic anti-tumor response

To determine whether the difference in tumor growth rates in recipient mice could be related to host immunity, we investigated immune cell populations in the spleen of mice by flow cytometry. There were significantly more neutrophils (CD45^+^CD11b^+^MHCII^-^Ly6G^+^Ly6^low^), monocytes (CD45^+^CD11b^+^MHCII^-^Ly6G^-^Ly6^high^), macrophages (CD45^+^CD11b^+^MHCII^+^), and dendritic cells (DCs) (CD45^+^CD11b^-^MHCII^+^CD11c^+^) in spleens from mice receiving the Pathogen CAG than in mice receiving the *Lachnospiraceae* CAG (**Fig. 5A**). In contrast, mice receiving the *Lachnospiraceae* CAG had multiple differences in the adaptive immune cell populations present in the spleen, including an increase in overall numbers of CD3+, CD4+ T cells, and cytotoxic CD8+ T cells, which is consistent with an active and effective anti-tumor immune response in these mice (**Fig. 5B**). There were also significantly more NK T cells (CD3^+^CD335^+^) in spleens of mice receiving the *Lachnospiraceae* CAG than in mice receiving the Pathogen CAG, with no change in NK cells (CD3^-^CD335^+^) (**Figs. 5B, S16**). These findings in the mouse model, together with the RNAseq analysis of human tumor tissue suggesting a strong and differential regulation of a microbiota-dependent immune response prompted us to characterize and quantify the immune infiltrate in the human biopsies using immunofluorescence. There was an increase in tumor-infiltrating CD15^+^ neutrophils in Pathogen CAG enriched tumors, while *Lachnospiraceae* CAG-enriched tumors had increased tumor-infiltrating total CD3^+^ T cells (**Figs. 5C-D, S17**). Together, these data demonstrate that a *Lachnospiraceae*-dominant microbiome is associated with a strong local and systemic adaptive immune response, while a Pathogen-enriched microbiome is associated with an immunosuppressive myeloid phenotype. Collectively these results suggest that the abundance of particular taxa is associated with specific host immune signatures that likely dictate tumor fate.

**Figure 5.**
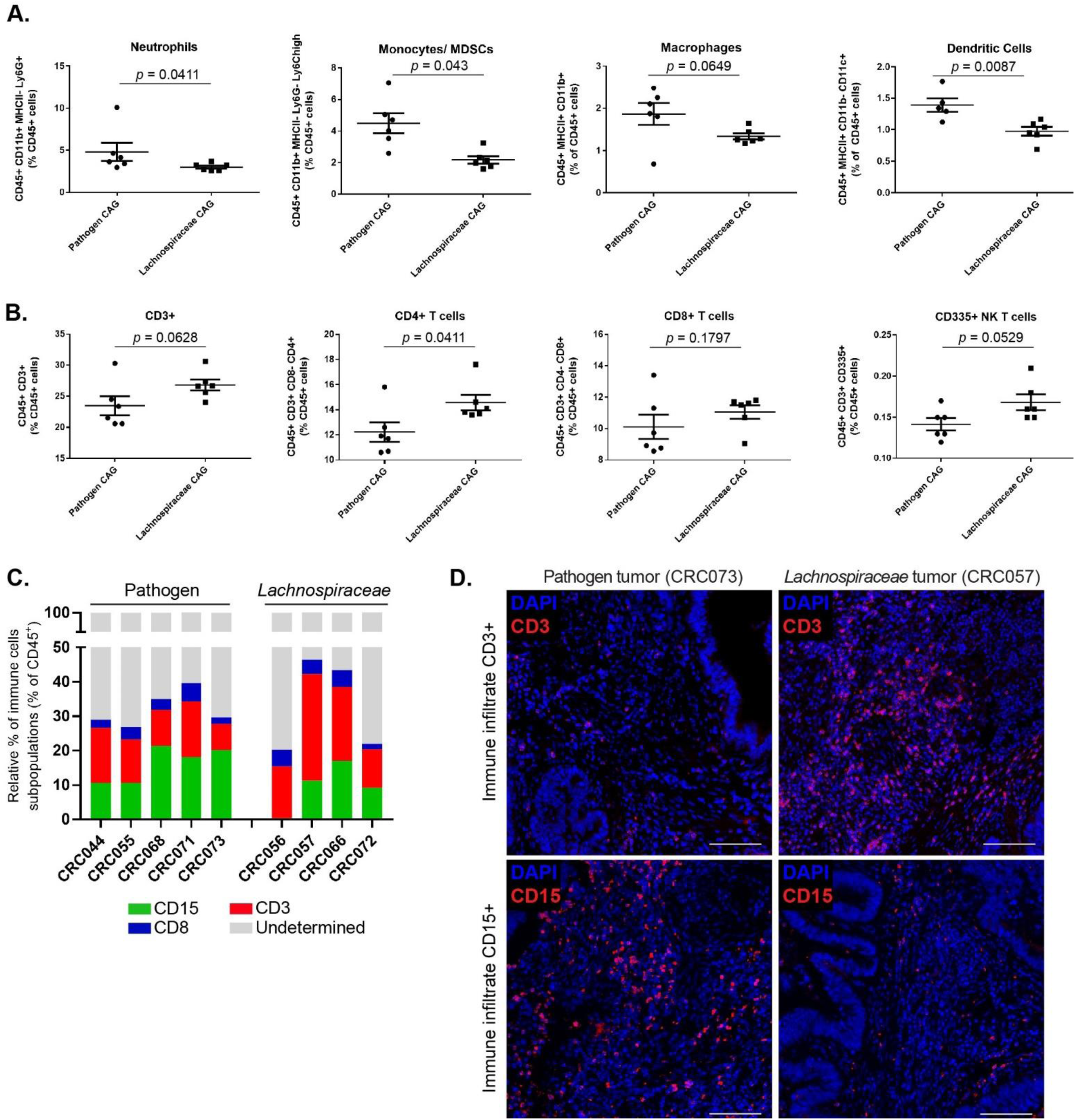
*Lachnospiraceae*-type microbiome colonization induces a strong immune infiltration and antitumoral immune response. **A.** Spleens from mice with the Pathogen CAG have more neutrophils (CD45^+^CD11b^+^MHCII^-^ Ly6G^+^Ly6^low^), monocytes (CD45^+^CD11b^+^MHCII^-^Ly6G^-^LyC^high^), macrophages (CD45^+^CD11b^+^MHCII^+^), and dendritic cells (CD45^+^MHCII^+^CD11b^-^CD11c^+^), as determined by flow cytometry gated on CD45^+^ cells. Panels show quantification of neutrophils, monocytes, macrophages, and dendritic cells. **B.** Spleens from mice with the *Lachnospiraceae* CAG have more CD3^+^, CD4^+^, CD8^+^, and NK T cells, as determined by flow cytometry gated on CD45^+^ cells. *P* values were determined by Mann-Whitney U test and are represented in each plot. Data indicate mean ±SEM. n= 6 biological replicates/group. **C.** Quantification of immune infiltrate (CD45^+^), T-cell infiltrate (CD3^+^ and CD8^+^) and neutrophil infiltrate (CD15^+^) in human CRC biopsies from Pathogen (n= 5) and *Lachnospiraceae*-enriched tumors (n= 4). 2 sections/tumor and 3 ROIs quantified per section (same ROIs were used to quantify different immune cell subpopulation in each tumor), means are shown, group comparison with one-way ANOVA. **D.** Immunofluorescence representative images of Pathogen CAG-enriched tumors (CRC073) and *Lachnospiraceae* CAG-enriched tumors (CRC057) human tumors showing that more CD3^+^ T cells (red) infiltrate into *Lachnospiraceae* tumors, while more CD15+ neutrophils infiltrate into Pathogen tumors. Counterstained with nuclear dye DAPI. All tumors were analyzed and for each tumor; 3 ROIs were quantified per section (n= 2 sections/tumor/staining). Scale bars 100µm.

## Discussion

Although human studies have suggested a causal link between an oncogenic microbiota and microbiota-induced immune response in CRC, there has been a lack of compelling causative evidence to support this concept. Our study consolidates the microbial ecology aspect of this thesis by showing that mice that received a human-derived *Lachnospiraceae* CAG microbiota developed much smaller orthotopic tumors than those receiving a Pathogen CAG microbiota. Based on our data and published studies we propose that the Pathogen-type microbiota, which is relatively over-abundant in CRC patients^4^, induces the recruitment of neutrophils, myeloid-derived suppressor cells, and M2 macrophages, enhancing tumor growth. Furthermore, our data suggest that the fiber-associated *Lachnospiraceae*-type microbiome has a protective role in the orthotopic model and induces a systemic T cell response, suggesting its contribution to a tumor-inhibiting microenvironment. These findings emphasize the important effects that microbiota abundance and composition can exert on immunomodulatory signals derived from the tumor microenvironment.

This study, in accordance with investigations of CRC patients, supports the existence of a cancer-specific signature^5, 35^. These putative “oncobacteria” include Clostridium XIVa pathobionts (*C. symbiosum*, *C. asparagiforme*, *C. hylemonae*, *C. bolteae,* and *C. hathewayi*)^39^, *Bacteroides sp*., including *B. fragilis*^40, 41^, as well as the high-sulfur-metabolizing *D. piger*^42^. In addition, previous studies in mice suggested that microbiota composition and structural organization contribute to CRC oncogenesis^43, 44^. However, many of the specific microbiota members associated with such responses vary substantially between studies^45^. For instance, none of the donors used in our study had detectable levels of *Fusobacterium nucleatum*, a known pro-carcinogenic bacterium that has been correlated with clinical outcome ^8, 46^. However, this is also consistent with studies where the presence of *F. nucleatum* in CRC biopsies largely varies between individuals^4^.

It is not currently clear whether the altered microbiome in CRC patients contributes to or is a consequence of disease. It may therefore be relevant that the pre-tumor (week 2) microbiota was predictive of tumor growth, which suggests that the pre-cancer microbiome may establish a permissive or protective environment for CRC development. Notably, here we identified a dominant human *Lachnospiraceae* microbiota as being protective in the mouse model. Specifically, commensals including *C. comes*, *R. lactaris*, *B. longum*, and *B. pseudocatenulatum* were largely responsible for this effect. *Bifidobacterium spp.* are known for their immunomodulatory effects^47^ and they were found to be depleted in CRC patients ^32, 48^, as well as in biofilm positive-inoculated mice^44^. Of note, the commensal *B. longum* has been reported to be enriched in patients with metastatic melanoma that respond to PD-1 immunotherapy and associated with improved immune-mediated tumor control^49^. This “protective” microbiome concept is consistent with the fact that the microbiome from mice that received the *Lachnospiraceae* CAG harbored a significantly higher abundance of health-related taxa groups^39^ (i.e. lost in disease), in the elderly and across multiple diseases, compared to the microbiome of mice that received the Pathogen CAG (**Fig. S18**).

The occurrence of genetic alterations involved in the initiation of human CRC has been suggested to be increased by a “dysbiotic” microbiota^6^. In that context, we suggest that it is relevant for CRC progression that the CAG microbiome types tested here contributed differently to the cancer hallmarks, including changes in the expression levels of genes involved in epithelial-mesenchymal transition, ^50, 51^, angiogenesis, and immune evasion. These pathways, which are associated with poor prognosis in CRC^50^, were overexpressed in patients harboring the Pathogen CAG. Interestingly, a recent study using a novel cancer mouse model with aberrant expression of the *Zeb2* gene, a master regulator of the epithelial-mesenchymal transition, showed that these mice were characterized by microbial dysbiosis and that the microbiota was necessary for the development of CRC in Zeb2-overexpressing mice^51^. It will now be interesting to investigate if the specific microbial components associate with consensus molecular subtypes^52^.

The composition and functionality of the immune infiltrate of the cancer patient are relevant for disease progression, metastasis, and immunotherapy treatment^53^. Abnormal immune responses, which are often accompanied by the recruitment of pro-inflammatory myeloid cells and overproduction of inflammatory cytokines, are associated with changes in the gut microbiota^6^. Specific bacterial species (e.g. *B. fragilis* and *P. anaerobius*) have been shown to trigger the secretion of chemokines that recruit immunosuppressive MDSCs, tumor-associated macrophages, and tumor-associated neutrophils^54, 55^. Albeit working in a pre-clinical model, we identified differences in the numbers of immune cells in the spleens of mice harboring the two different microbiota types, which is consistent with the transcriptomic differences observed in the tumor-immune genes of this cohort of CRC patients. Manipulation of the microbiota with the goal to modulate towards a “cold” tumor, characteristic of patients with sporadic CRC, towards a “hot” tumor environment, which is characterized by a high infiltration of T cells and an effective response to checkpoint blockade, is a promising and valid therapeutic avenue, especially when in combination with current immunotherapies.

Although we compared our findings in the mouse model to six large human CRC studies and confirmed many of the identified key taxa, we acknowledge the limitation of the use of only one representative donor for each microbiota type. Another limitation in this study is the fact that it is not powered to detect very small differences in human genes between the two different CAGs due to the small number of patient samples analyzed, which limits our ability to detect interactions between specific driver mutations and microbiome structure. Moreover, whether the microbiota-induced immunity affects the mutation burden of the tumor or vice-versa, remains to be determined.

In this study, we expanded on previous work by us and others to identify specific taxa and respective metabolites that correlate with CRC development and to suggest a mechanism that involves immune modulation by the microbiota and its metabolites. Future studies involving the culture of these microbial taxa, comparative genomics to evaluate strain-specificity and variability, and *in vitro* functional assays with defined bacterial consortia and/or their metabolites will be essential to understand if (and which) bacterial compounds can ultimately be used as immunomodulators in CRC.

## Acknowledgments

We thank the APC Germ-Free facility staff (Frances O’Brien and Tara O’Driscoll), APC Flow Cytometry manager Pepi Stamou; APC research nurses (Orlagh O’Connor, Joanne Carroll, and Ciara Tobin); Juliet Barry at Cork Cancer Research Centre (CCRC) for processing histology samples; and Jan Soetaert (QMUL) for assistance on image analysis. Special thanks to all patients that generously participated in this study. The graphical abstract and figures 1A and 2B were created with Biorender.com

## Author contributions

ASA, FS and PWOT designed the study; ASA, TTT, TSG, CR, and BF performed microbiota data analysis; CF performed RNAseq analysis on human samples; ASA, LAA, PP, WF, CMH and RJ performed experiments; ASA, RJ and IS performed immune infiltrate data analysis; MOR was responsible for patient sample acquisition; ASA and PWOT wrote the manuscript and obtained funding for the study.

## Materials and Methods

### Patients cohort and sample collection

All clinical studies were conducted after informed consent of the patients, following the guidelines of the Declaration of Helsinki. Ethical approval for this study was granted by the Clinical Research Ethics Committee of the University College Cork, under the study number APC033. Patients’ data were anonymized and stored under European Union General Data Protection Regulation.

Detailed clinical and pathological information on the patients is presented in **Table S1**. A total of 32 treatment-naïve patients with adenomas or CRC were included in this study. Exclusion criteria included a personal history of CRC, inflammatory bowel disease (IBD), or inflammatory bowel syndrome (IBS), as well as chemotherapy or radiation therapy treatments. Individuals were not treated with antibiotics in the month prior to surgery but were administered antibiotics intravenously during surgery. Dietary data for each patient were collected using a validated Food Frequency Questionnaire (FFQ)^22^. Control subjects (i.e. routine colonoscopy) included individuals without a history of CRC, IBD or IBS, or antibiotic usage within 3 months, described elsewhere^4^.

Resection samples were collected from CRC patients undergoing surgery at Mercy University Hospital, Cork. Samples were collected from the tumor site (ON), 2-3 cm far from the tumor margin (OFF), and paired healthy tissue (approximately 10 cm from the tumor site). Samples collected were rapidly preserved in 1) RNA later for sequencing purposes; 2) methacarn for histology and 3) snap-frozen for subsequent analysis. Bowel preparations before surgery or colonoscopy were determined by the surgeon and are detailed in **Table S1**.

### DNA and RNA extraction of human biopsies

Human colon resections were placed in RNAlater (Qiagen) at the time of resection, stored at 4°C for 12 h, and then stored at −20°C until processing. Genomic DNA and total RNA were extracted using the AllPrep DNA/RNA kit (Qiagen). For tissue samples, ∼20 mg of tissue was placed into bead tubes containing 250 µl of 0.1 mm sterile glass beads (Biospec Products) and three 3–4 mm sterile glass beads (Biospec Products). Next, 600 mL of buffer RLT (Qiagen) containing 1% β-mercaptoethanol was added and the sample was homogenized in a MagnaLyzer (Roche) for two pulses of 15 s each at full speed. The extraction was then carried out using the AllPrep DNA/RNA extraction kit (Qiagen), following the manufacturer’s instructions. Genomic DNA was quantified using the Nanodrop 1000 (Thermo Scientific) and total RNA was quantified using the Bioanalyser (Agilent).

### Human fecal inoculum preparation

Two human donors were selected from the CRC patients’ cohort based on their mucosal microbiota compositional profiles. Donor 1 (CRC044) is a female, 66 years old, diagnosed with a T3N2b rectum adenocarcinoma; donor 2 (CRC056) is a male, 65 years old, diagnosed with a tubulovillous adenoma with low-grade dysplasia (**Table 1 and Figure 2A**). Stool samples were transferred to an anaerobic chamber immediately after voiding, transported to the lab in anaerobic bags, and transferred to an anaerobic hood in less than an hour. Fresh stools were diluted (ratio 1:10) in sterile pre-reduced PBS with 20% glycerol and stored at − 80°C in aliquots until further use.

### DNA extraction of human fecal samples

Genomic DNA was extracted from fecal samples following the Repeated Beat Beating (RBB) Method^56^, with the following modifications. Samples (0.25 g) were placed in sterile tubes containing one 3.5 mm zirconia bead and one scoop of 0.1 mm and 1 mm beads, respectively (Thistle Scientific, UK). Fecal samples were homogenized via bead beating for 60 seconds (Mini-Beadbeater™, BioSpec Products), with the samples cooled on ice for 60 seconds before another 60 seconds bead beating. Samples were then incubated at 70°C for 15 min and centrifuged at full speed for 5 min. Pooled supernatants were incubated with 350 ml of 7.5 M ammonium acetate (Sigma) and incubate on ice for 5 min. The remaining steps of DNA purification were performed using QIAamp columns (Qiagen). Genomic DNA was quantified using the Nanodrop 1000 (Thermo Scientific). Extracted genomic DNA was stored at −20°C until amplification.

### MC-38 cells culture

Murine C57BL/6 MC-38 colorectal tumor cells were obtained from Kerafast (ENH204-FP) and maintained in 5% CO_2_ at 37°C in Dulbecco’s modified MEM (DMEM) medium supplemented with 10% heat-inactivated fetal bovine serum (Sigma), 2mM L-glutamine, 0.1 mM nonessential amino acids, 1 mM sodium pyruvate, 10mM HEPES and 100 units/ml penicillin/streptomycin antibiotic solution (all reagents from Gibco-Invitrogen). Cells were tested for Mycoplasma contamination every 4 - 6 weeks and before each experiment (Mycoalert Mycoplasma Detection kit, Lonza).

### Germ-free MC-38 mouse model

All animal protocols were approved by the Animal Experimentation Ethics Committee at University College Cork and by the Health Products Regulatory Authority (HPRA) of Ireland, per EU Directive 2010/63/EU (HPRA Project authorization number AE19130/P055).

Germ-free (GF) mice were bred and maintained at the APC Germ-Free facility in dedicated axenic isolators (Bell Isolation Systems). Germ-free status was routinely monitored by culture-dependent methods. Age-matched male C57BL/6 GF mice, 6-10 weeks of age, were group-housed 3-4 and transferred into sterile individual ventilated cages (IVCs) (Arrowmight, Hereford, UK) before undergoing human microbiota administration. Mice were kept in a 12-hour light-dark cycle and on ad libitum diet RM1 (autoclaved) (Special Diet Services, #0103). Water and diet were batched at the beginning of the experiment to exclude possible variations between batches. An overview of the experimental study design is presented in **Figure 2B**. Animals were pipette-dosed with 100 µl of fecal slurry or control PBS, per day for 3 consecutive days. Groups were as follows: “Pathogen CAG” group, inoculated with fecal slurry from donor 1 (patient CRC044); “*Lachnospiraceae* CAG” group, inoculated with fecal slurry from donor 2 (patient CRC056); and, GF control group inoculated with 100 µl of reduced PBS + glycerol 20%. After microbiota administration and colonization, MC-38 cells were orthotopically injected into the rectal submucosa, as previously described^27^. Briefly, mice were anesthetized using a mixture of Ketamine/Medetomidine (75 mg/kg ketamine (100mg/ml), 0.5 mg/kg medetomidine (1mg/ml) subcutaneously. Injection of 20 µl of 5×10^5^ (experiment1) or 1×10^5^ (experiment 2) MC-38 cancer cells was performed using an insulin syringe on the right flank. Fecal and blood samples were collected from each animal at various time points (weeks 0 to 5) as indicated in **Figure 2B**. Two to three fecal pellets were collected at each time point and were immediately frozen in dry ice before being transferred to −80°C. Tumor size was measured by caliper at endpoint (week 4 in experiment 1 and week 5 in experiment 2), and tumor volume was calculated as (length × width^2^)/2.

### Genomic DNA extraction and microbiota profiling of murine fecal samples

Total DNA was extracted from mouse fecal samples using QIAamp DNeasy Blood and Tissue Kit (Qiagen, UK) according to the manufacturer’s protocol and as previously described^57^. Genomic DNA was quantified using the Nanodrop 1000 (Thermo Scientific, Ireland). Extracted genomic DNA was stored at −20°C until amplification.

### Sequencing, Taxonomic and Functional Profiling

#### 16S rRNA Gene Amplicon Sequencing

The V3-V4 region of the 16S rRNA gene was amplified, sequenced, and analyzed as described before^58^. Amplification was performed with the universal 16S rRNA gene primer pair S-D-Bact-0341-b-S-17 and S-D-Bact-0785-a-A-21^59^. The Phusion High-Fidelity PCR Master Mix (ThermoFisher Scientific, USA) was used for the amplification. The sequencing library was prepared using the Nextera XT V.2 Index Kit (Sets A and D, Illumina) according to the Illumina 16S MiSeq Sequencing Library protocol. The PCR products were purified with the SPRIselect reagent kit (Beckman Coulter, Inc., USA). Amplicons were quantified with a Qubit dsDNA HS Assay Kit (Thermo Fischer Scientific) and pooled at the same concentration. Sequencing was performed on an Illumina MiSeq Platform (2×250 bp reads for human samples and 2×300 bp reads for mouse samples) by the Teagasc Next Generation DNA Sequencing Facility (Fermoy, Ireland).

#### Microbiota composition analysis of 16S rRNA amplicon sequencing data

Primers were removed from raw sequences using Trimmomatic^60^ (v0.36). Paired-end sequencing reads (2×250 bp or 2×300 bp) were joined using FLASH^61^ (v1.2.8). Demultiplexing and quality filtering were performed using the QIIME package^62^ (v1.9.1). The USEARCH^63^ sequence analysis tool (v8.1.186) was applied for further quality filtering and *de novo* clustering to form operational taxonomic units (OTUs). The sequences were initially filtered by length and sorted by size, and single unique sequences were removed. The remaining sequences were clustered into OTUs at 97% similarity. Subsequently, chimeras were removed with UCHIME^64^, using the GOLD reference database. The original quality-filtered sequences were mapped against the OTUs at 97% sequence identity. OTU representative sequences were classified with a confidence threshold of 80% to taxonomic ranks from phylum to genus level by mothur (v1.36.1) using the RDP reference database (trainset 16^65^) and to species level by SPINGO^66^ (v1.3) using the RDP reference database (v11.4). Alpha (α) and beta (β) diversity analyses were performed in QIIME on a rarefied OTU table. The sequences were aligned using the PyNast tool^67^ in QIIme to generate α-diversity indices (Shannon, Simpson, PD whole tree, Chao1, and Observed Species), and β-diversity indices (Bray-Curtis, Weighted UniFrac, and Unweighted UniFrac).

### Shallow shotgun analysis

#### Pre-processing of shotgun sequence data

Pre-processing of raw shotgun sequence reads was performed using a similar approach adopted by previous studies^68^. To summarize, the reads were quality trimmed using Trimmomatic (v0.39, with default parameters)^60^; followed by removing reads originating from the host genome bowtie2 (v2.3.4 with default parameters) (with *Mus musculus* genome version 9 for mouse fecal metagenomes and human genome hg38 for donor fecal metagenomes)^69^.

#### Taxonomic, functional, and strain-wise profiling

The taxonomic and functional profiling of the metagenomes was performed using the HUMAnN2 pipeline with the clade-specific marker gene-based metaphlan2 as the taxonomic classifier^70, 71^. Pathway and gene-family abundances obtained using the UniProt mapping scheme were subsequently converted into the KEGG-specific mapping scheme using the legacy databases of humann2 (as described in Keohane *et al.*^68^). Strain-wise variations were profiled using Strainphlan2^72^. Inferred metabolite production and consumption profiles were obtained using a similar approach as adopted in Ghosh *et al.*^39^.

### RNA-Sequencing

RNA sequencing libraries were prepared by Genewiz with the Standard RNA-seq protocol for tumor samples and by GATC for healthy control samples. Tumor samples were sequenced on an Illumina HiSeq instrument with 150-bp paired-end reads to an average depth of 29 million pairs of reads per sample. Healthy control samples were sequenced with 100-bp paired-end reads to an average depth of 45 million pairs of reads per sample.

### RNAseq transcriptome analysis

FastQC software (v0.11.5) was performed to assess the quality control checks of paired-end sequencing reads. The TrimGalore (v0.6.5) tool was used with Cutadapt (v1.15) and FastQC to apply quality and adapter trimming to FASTQ files. STAR (v2.7.3a) was used to align trimmed reads to the human genome (*Homo sapiens* high coverage assembly GRCh38 from the Genome Reference Consortium – GRCh38.p13) with the --quantMode GeneCounts option to output read counts per gene. The Bioconductor package EdgeR (v3.28.1) was applied in R (v3.6.3) to identify statistically significant differentially expressed genes between patient groups. Biological and technical variation was accounted for by the negative binomial distribution of RNAseq count data using a generalization of the Poisson distribution model. The filterByExpr function was applied to remove lowly expressed genes. The data was normalized across library sizes, between samples using the trimmed mean of M-values (TMM) normalization method. Tagwise dispersions were estimated for the normalized dataset. P values from multiple comparisons were corrected with the Benjamini-Hochberg method in EdgeR. For the comparisons between tumors and healthy controls, genes were considered significantly differentially expressed with an FDR adjusted *p* value < 0.1. For the comparisons between Pathogen and *Lachnospiraceae* enriched tumor genes were considered significantly differentially expressed with a *p* value < 0.05. Voom “variance modeling at the observational level” method within edgeR was used to output normalized read counts as LogCPM values. These were used to perform hierarchical clustering and to construct heatmaps in Gene Pattern’s online server (v3.9.11), volcano plots in Gene Patterns Multiplot Studio (v1.5.2), to estimate the abundances of immune cell types in a mixed cell population with CIBERSORTx^25, 26^ signature genes (LM22), and to perform Gene Set Enrichment Analysis (GSEA) (v4.1.0) with annotated HALLMARK genesets from the MSigDB (Molecular Signatures Database) collections (v6.2). Venn diagrams were constructed using InteractiVenn. Further statistical analysis of estimated abundances of immune cell types from CIBERSORTx involved students t-tests between patient groups within GraphPad Prism (v6). Summary statistics for the RNA Seq data analysis is given in **Table S3 and S4**.

### Statistical analysis

Statistical analysis, data visualization and machine learning-based analyses were carried out in R statistical software package (v3.4.0). Principal Coordinate Analyses (PCoA) were performed using the dudi.pco function of the ade4 package (v1.7-15). Two-dimensional PCoA plots were created using the ggplot2 package (v2.2.1). Permutational multivariate analysis of variance (PERMANOVA) analyses to test for statistical difference in β-diversity, the gut microbiome profiles as well as the strain variations of the different species were performed using the adonis function from the vegan package (v2.5-6). Spearman Distances of the species abundances across samples and the species-specific strain-wise distances were provided as inputs to the adonis function. PERMANOVA analysis of the strain-wise variations for each species (obtained using Strainphlan) was performed in a time-point specific manner (separately for week 2 and week 5), after adjusting for the donor and abundance of the given species as confounders. Effect size calculations (Cohen’s D) were performed using the effsize package (v0.8.1). We excluded from the analysis any bacterial taxa or metabolites that were detected in less than 50% of the samples of each group. Significant variations in α-diversity, taxa/gene relative abundance, metabolites abundance, and pathway coverages were assessed on median values of the technical replicates using the Mann-Whitney U test for unpaired data or Wilcoxon signed-rank test for paired data. The Kruskal-Wallis test followed by Dunn’s post hoc test with Benjamini-Hochberg *p* value adjustment for multiple testing was applied when comparing more than two experimental groups. The bar plots showing different taxonomic level classification were created using the ggplot2 package. Taxa below 1% sample abundance and the unclassified taxa were grouped into the “Other” category. *P* values from multiple comparisons were adjusted for the FDR using the Benjamini-Hochberg method (implemented in the p.adjust R function). Significance was assumed for adjusted p values ≤0.05, if not stated otherwise. Correlations between metabolite and taxa relative abundances were calculated using standard Spearman’s rank correlation using the ‘corr’ function implemented in the ‘psych’ module of R and hierarchical clustering was computed using the hclust function in R (method “complete”). Features (that is taxa at various time-points and inferred metabolites) that showed significant Spearman correlations with tumor volumes (FDR < 0.1, obtained after *p* value adjustment using the Benjamini-Hochberg method) were visualized using the ggplot2 package.

### Machine-learning based analysis

For comparative validation on a global scale, we utilized the curatedMetagenomicData repository to six additional case-control datasets containing human fecal shotgun metagenome data from greater than 600 individuals consisting of CRC patients and controls^73^ (this was referred to as the ‘Global Reference CRC cohort’^32–37)^. For comparing taxa abundances or inferred metabolite inferences within this Global Reference CRC cohort, we adopted a two-step procedure. First, for inter-dataset variability in the detection of various taxa, we performed across sample rank normalizations of taxa abundances separately within each dataset corresponding to the six studies. This limited the abundance range of each taxon from 0 to 1 uniformly for all the six studies. Subsequently, the rank normalized profiles for the six studies were combined and the comparison of rank normalized taxa abundances were compared between CRC patients and non-diseased individuals using Mann-Whitney U tests. Machine-learning based analyses consisting of evaluating the disease predictive ability of various markers in the Global Reference CRC cohort as well as within our dataset were performed using Random Forest approach (using the randomForest module implemented the R-programming interface). Iterative random forest classifiers built by taking repeated 50% subsets of ‘test’ and ‘training’ samples were obtained using the same methodology as used in Ghosh *et al*.^39^.

### Flow cytometry

Spleens were harvested 19 days after MC-38 cancer cells injection (Experiment 2) and processed for flow cytometry analysis as previously described^74^. For staining, single cells (1×10^6^) were pre-incubated with purified anti-mouse CD16/CD32 (clone 2.4G2, Biolegend) for 10 min on ice, and then stained with the appropriate surface markers antibodies at 4°C for 30 min in the dark. Zombie Red (BioLegend) was used to differentiate between dead and live cells. The stained populations were analyzed using a BD FACSCelesta™ flow cytometer (BD, USA) and FlowJo software (BD, v10). Antibodies were titrated for optimal staining. Antibodies used were: CD45-BV510 (Clone 30-F11, dil 1/100), CD274 (B7-H1, PD-L1)-BV605 (Clone 10F.9G2, dil 1/100), Ly6G-BV786 (clone 1A8, dil 1/100), CD11b-FITC (clone M1/70, dil 1/100), CD103-PE (Clone 2E7, dil 1/200), F4/80-PerCP-Cy5.5 (Clone BM9, dil 1/80), CD11c-APC (Clone N418, dil 1/200), Ly6C-AF700 (clone HK1.4, dil 1/200), CD3-FITC (Clone 17A2, dil 1/100), NKp46-PE (clone 29A1.4, dil 1/50), CD4-PerCP-Cy5.5 (Clone RM4-4, dil 1/200), CD279 (PD-1)-APC (clone 29F.1A12, dil 1/200), CD8-AF700 (Clone 53-6.7, dil 1/200), all from BioLegend; MHC II (I-A/I-E)-APC-eFluor 780 (Clone M5/114.15.2, dil 1/200) from eBioscience.

### Immunofluorescence staining, imaging, and quantification

Methacarn-fixed paraffin sections of human CRCs were blocked with 10% fetal bovine serum, 2% BSA, 0.02% fish skin gelatin, 0.05% TritonX100 (Sigma) and 0.05% Tween (Sigma). After 1h blocking at room temperature, primary antibodies were incubated overnight at 4°C, followed by 2h incubation at room temperature in secondary antibody.

The following primary antibodies were used: mouse anti-CD45 (Leica Biosystems, NCL-L-LCA, 1/70), mouse anti-CD3 (Leica Biosystems CD3-565-L-CE, 1/100), mouse anti-CD8 (Novus Biologicals, NBP2-32952, 1/200), mouse anti-CD15 (Cell Signalling, SSEA1 MC480 #4744S, 1/1500). AlexaFluor555 donkey anti-mouse (all Invitrogen, purchased from ThermoFisher Scientific, 1/300) as use used as secondary antibody. DAPI (Life Technologies) was used as a nuclear counterstain. Slides were mounted using ProLong Gold anti-fade reagent (Life Technologies). TissueFAXS Quantitative Imaging System (TissueGnostics, Vienna, Austria) was used to acquire images from the slides. The TissueFAXS uses a standard widefield epi-fluorescence Zeiss AXIO Observer.Z1 Inverted Microscope (with high efficiency fluorochrome specific DAPI, GFP, CFP, Cy3, and Cy5). Images were captured with a Hamamatsu ORCA-Flash 4.0 CMOS Camera. The entire slide (75 × 25 mm^2^) was scanned at low magnification using a 5× objective to identify the location of the tissue on the slide, followed by acquisition in multiple sequential tiles at 20× high magnification used for all downstream analysis. Image processing and analysis was performed using StrataQuest software version 6.0.1.209 (TissueGnostics, Vienna, Austria). Image processing included reconstruction of whole images and creation of *in silico* multiplexed images. Tissue cytometry and backgating into the tissue images were used for quantitation and visualization of the *in silico* data. In brief, several algorithms were used: to isolate cells by DAPI staining, to create a ring mask and identify non-nuclear staining starting from the centroid of the identified nucleus and stopping at the exterior of the biomarker, to identify the biomarker for the cell phenotype (CD45, CD3, CD8, CD15 stainings), allowing isolation of individual cells by a specific phenotype. Global standard measurements were computed for area (μm^2^), mean fluorescence intensity, perimeter (μm), compactness, and cell location (Cartesian coordinates). 2D dot scatterplots were created for each ROI (region of interest) containing the mean fluorescence intensity of one biomarker on each axis. Using the backgating algorithm, threshold cutoffs were manually positioned to include or exclude cell subpopulations. Total number of cells (DAPI^+^) and positively stained cells for CD45, CD3, CD8, and CD15 were segmented in each ROI.

### Statistical analysis and reproducibility

Statistics for 16S rRNA gene sequencing, shotgun metagenomic sequencing, and human RNA-seq are described above. Other data were plotted and analyzed using GraphPad Prism 7 for Windows, and specific statistical methods used are indicated in the text or figure legends. Briefly, a two-tailed Mann-Whitney U test was used for comparison between two independent groups. For multi-group comparisons, one-way or two-way ANOVA followed by Dunn’s multiple comparisons test was performed. Only statistically significant differences are indicated in the figures. Exact *p* values and statistical tests used for each panel are reported in the source data. The number of samples analyzed in each group are indicated in each figure.

Sample size of mice follows the 3 Rs (replace, reduce, and refine). Mice were randomly assigned to experimental groups and matched to the best age. No data were excluded from the analyses and details on experiment repetition are given in the respective figure legends. The investigators were not blinded to group allocation for mouse stool collection and euthanasia to avoid sample cross-contamination.

