## Supplementary material for "Fiber-associated *Lachnospiraceae* reduce colon tumorigenesis by modulation of the tumor-immune microenvironment": Fig S

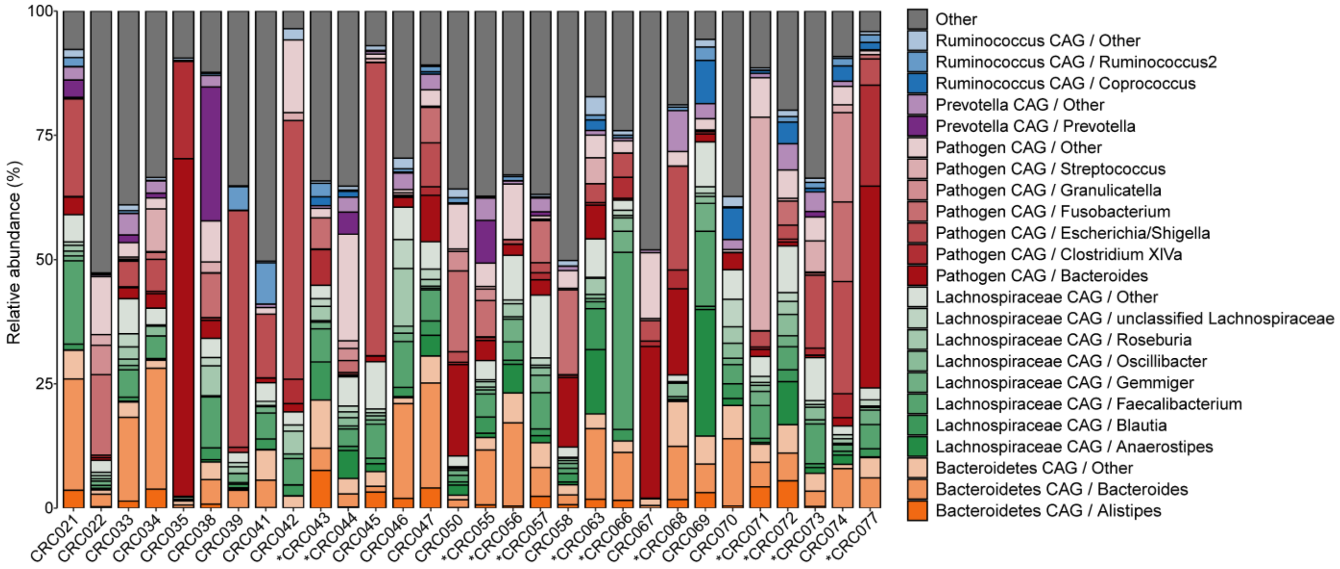


**Figure S1**. Human microbiota composition presented as proportional abundance of bacterial CAGs in patients with adenomas or CRC. Only genera whose mean relative abundances in all samples was >1% are presented. Low abundance genera of each CAG group are combined to “CAG/Other” group, whereas “Other” are taxa not included in any CAG groups. Stars (*) indicate the 12 patients selected for RNAseq analysis.

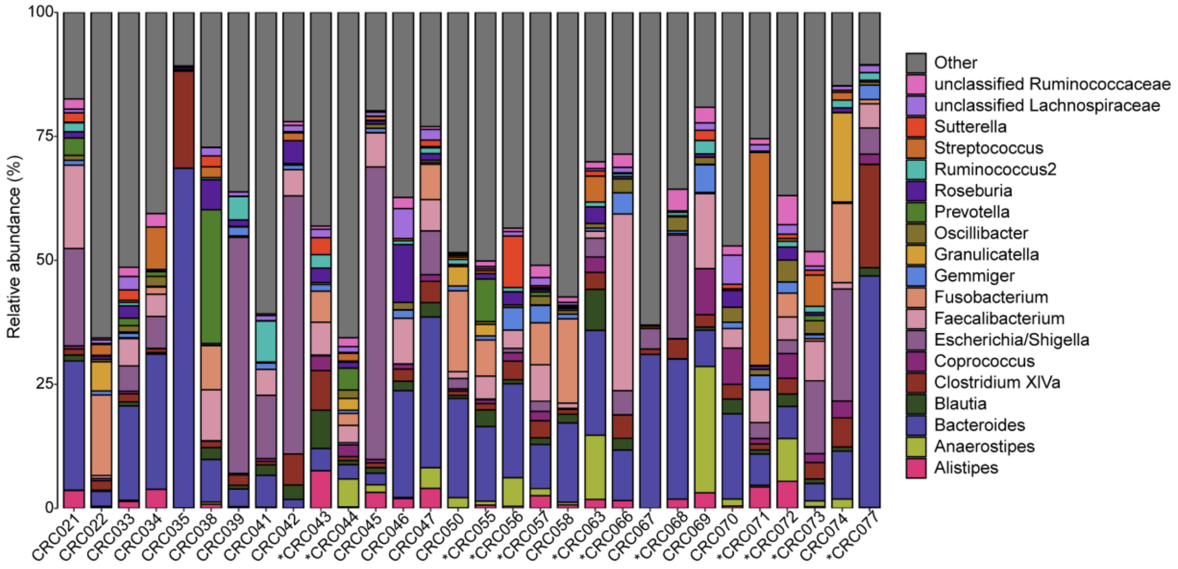


**Figure S2** . Genus level microbiota composition of patient biopsies. Only bacterial genera that were present at >1% relative abundance are presented. Stars (*) indicate the 12 patients selected for RNAseq analysis.

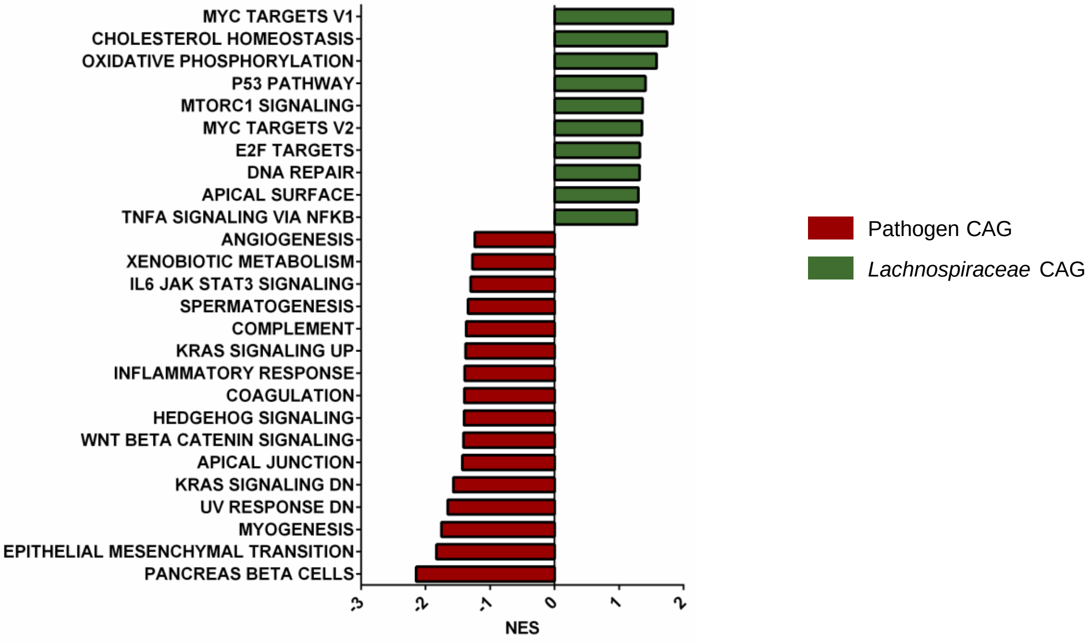


**Figure S3.** Gene Set Enrichment Analysis (GSEA) highlights the different transcriptome of Pathogen CAG (n= 6) and *Lachnospiraceae* (n= 6) CAG tumors. Gene Set Enrichment Analysis (GSEA) displaying significantly (FDR<0.25) enriched Hallmark gene sets (MSigDB) ranked based on normalized expression score (NES). The histogram highlights pathways uniquely enriched in Pathogen CAG and *Lachnospiraceae* CAG-enriched tumors compared to healthy controls (n= 10).

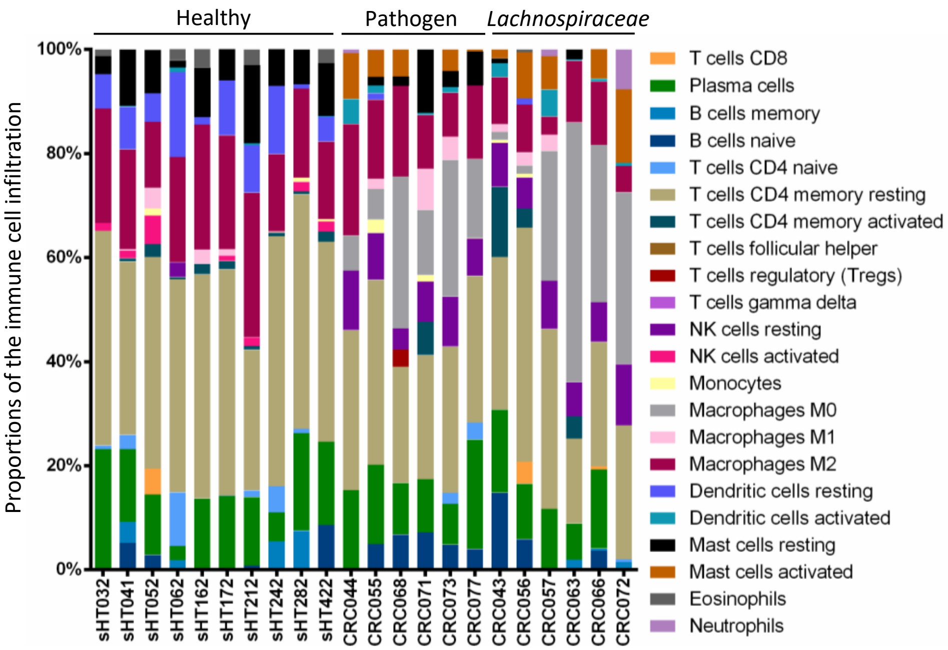


**Figure S4.** Landscape of immune infiltration in healthy controls, Pathogen CAG, or *Lachnospiraceae* CAG-enriched tumors. Bar charts of 22 immune cell proportions as calculated using CIBERSORTx using the LM22 signature matrix file.

**A.**

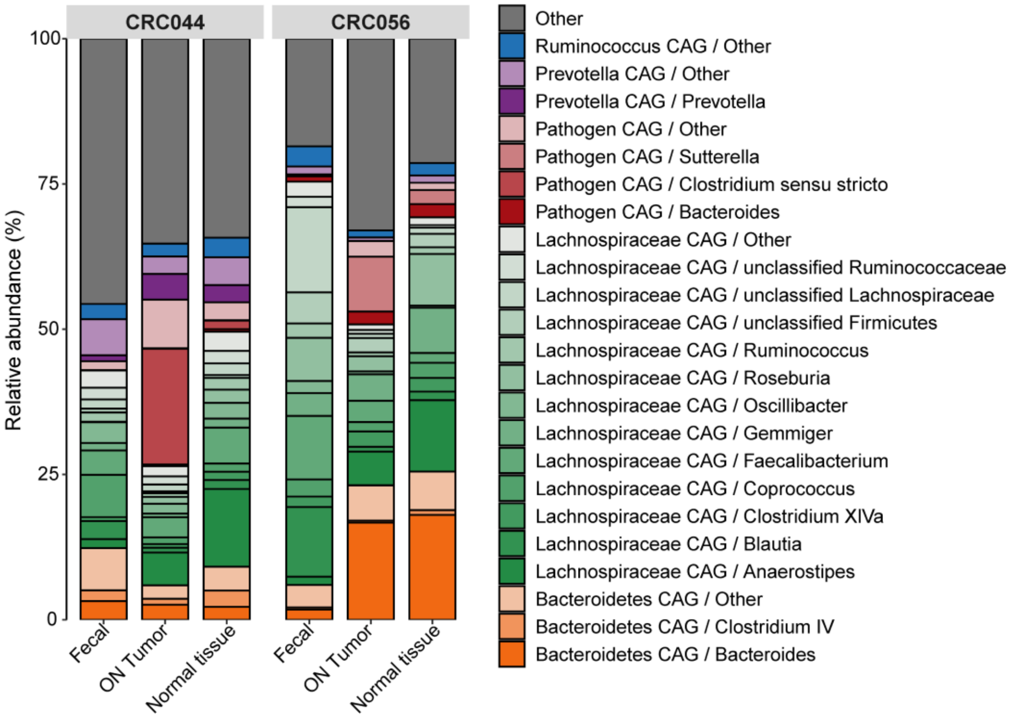


**B.**

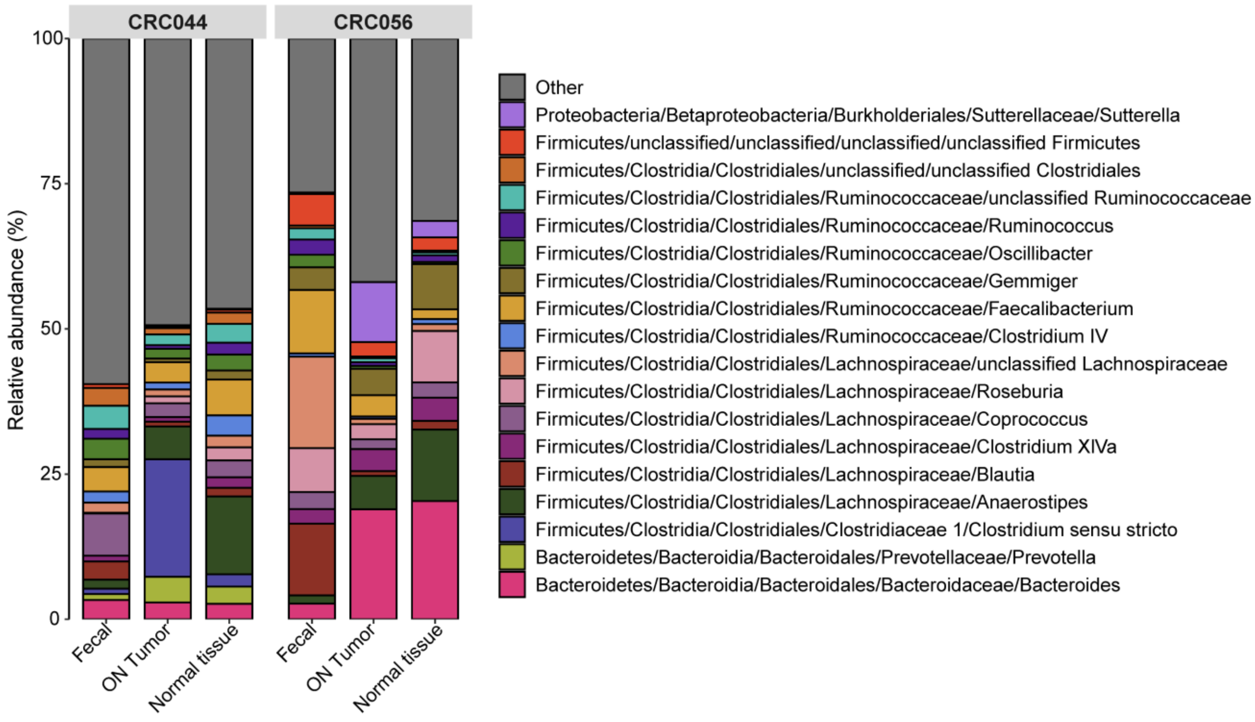


**Fig. S5. A.** Human microbiota composition at genus level of each bacterial CAG of fecal, ON tumor, and paired normal tissue of donors CRC044 (Pathogen CAG) and CRC056 (*Lachnospiraceae* CAG). **B.** Human microbiota composition at genus level of fecal, ON tumor, and paired normal tissue of each donor. Bacterial genera at >1% relative abundance are presented.

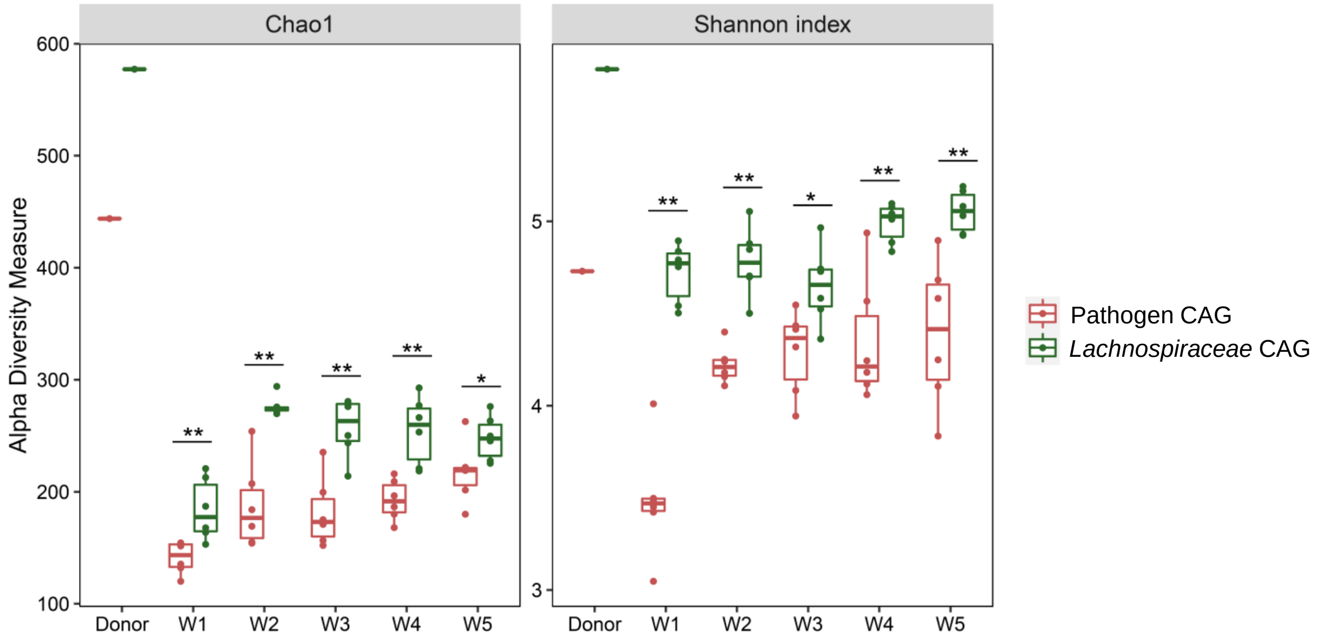


**Fig S6.** Alpha-diversity indexes (Chao1 and Shannon) of Pathogen CAG versus *Lachnospiraceae* CAG fecal microbiomes. Bar plots show the alpha-diversity indexes of the human donors and murine microbiota composition at weeks (W) 1-5. *p* values were calculated with Mann-Whitney U test. *$p \leq 0.05$, **$p \leq 0.01$.

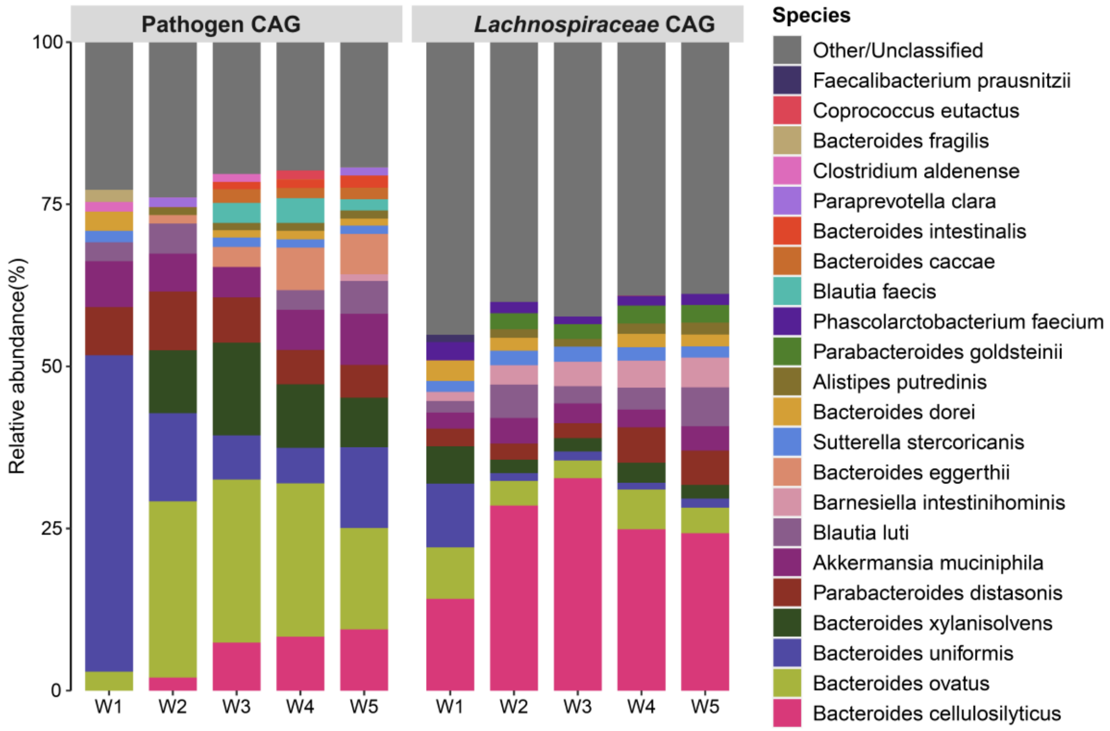


**Fig S7.** Fecal microbiota composition profiles at species level in mice that received Pathogen CAG and *Lachnospiraceae* CAG over time (weeks 1-5) revealed by 16S sequencing. Only classified species that were present at >1% relative abundance in any of the time points are presented. n= 6 mice per experimental group.

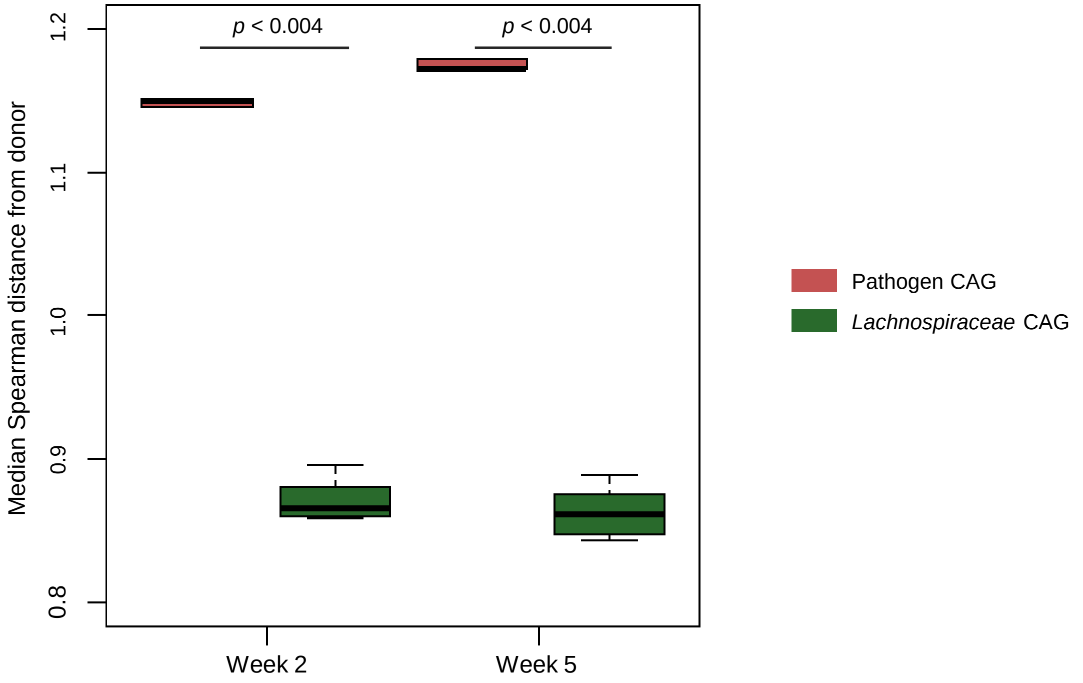


**Fig S8.** Boxplots showing the distribution of the Spearman distances of the different mice fecal microbiomes from that of their corresponding donors at week 2 and week 5. *p* values were calculated with Mann-Whitney U test.

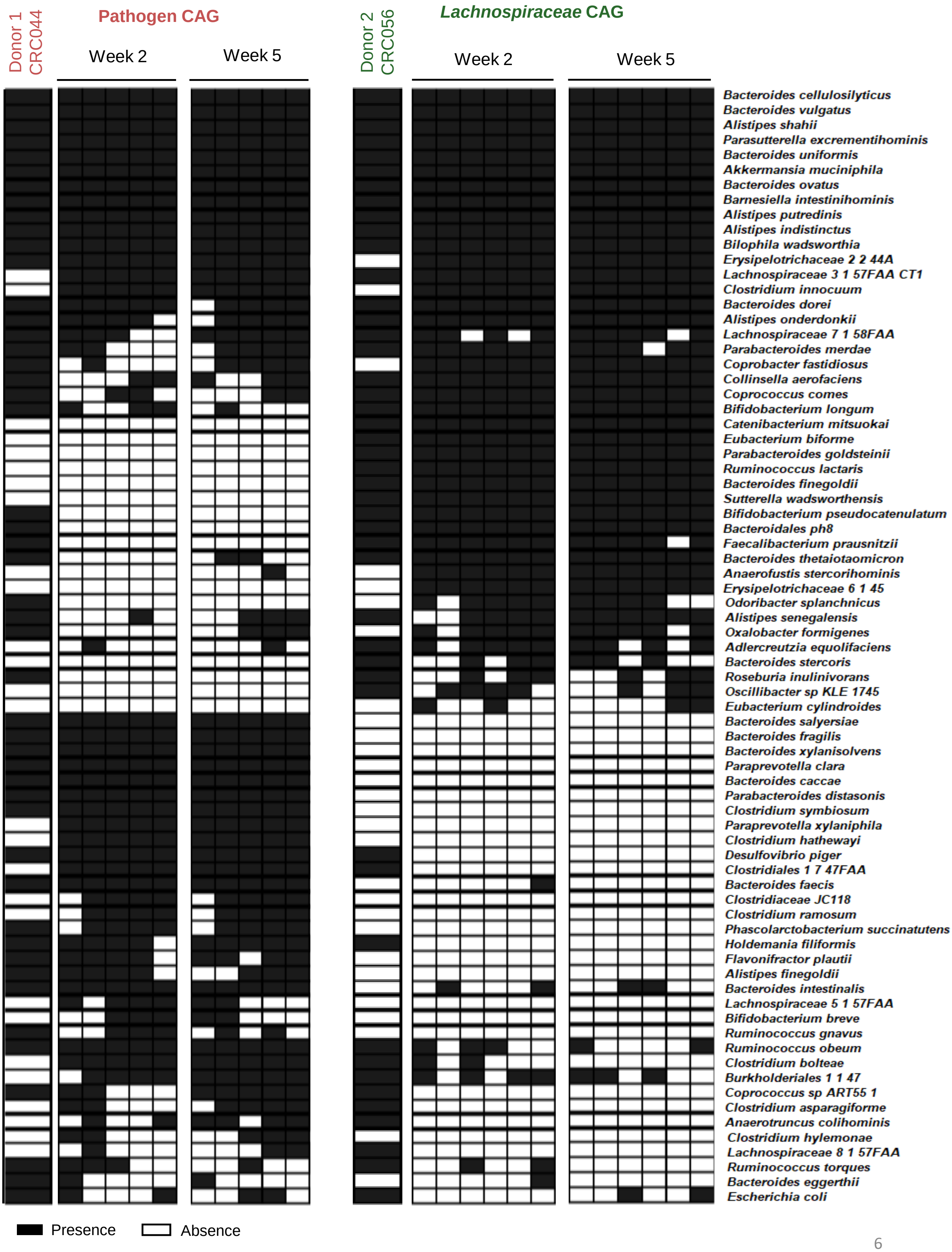




**Fig S9** Shared and unique bacterial species of the two human donors and murine microbiota composition of the Pathogen and *Lachnospiraceae* CAGs at weeks 2 and 5, identified by shallow shotgun sequencing analysis. Presence is color-coded in black and absence is color-coded in white.

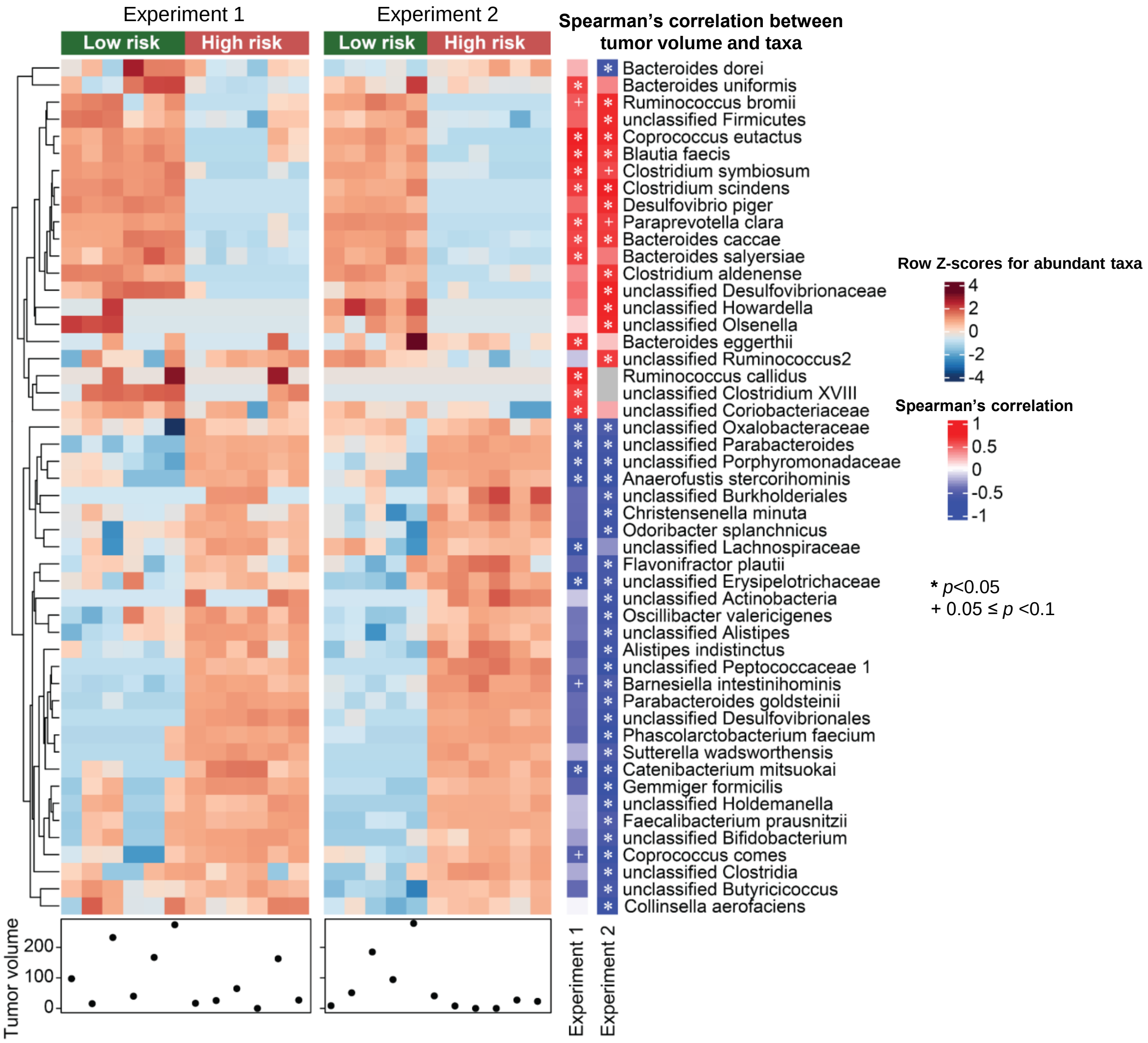


**Fig S10.** Heatmap and hierarchical clustering showing the relative abundance of several bacterial species associated with tumor volume. The right heatmap shows Spearman correlation between species and tumor volume for experiment 1 and experiment 2, red color indicates positive correlation and blue color indicates negative correlation. Only correlation coefficients value greater than 0.5 are shown, and significance correlation is given as nominal *p* values: $*p < 0.05$, $+\ 0.05 \leq p < 0.1$. The left heatmap shows the relative abundance (log 10 transformation) of the tumor-correlated species across the samples in each experiment. Color intensity of the heatmap is based on row z-scores of the relative abundance, ranging from blue (low abundance) to red (high abundance). The dendrogram at the left represents hierarchical clustering of bacterial communities based on the Euclidean distance. Each column represents an individual mouse with tumor volume ($mm^3$) shown by the bottom plot.

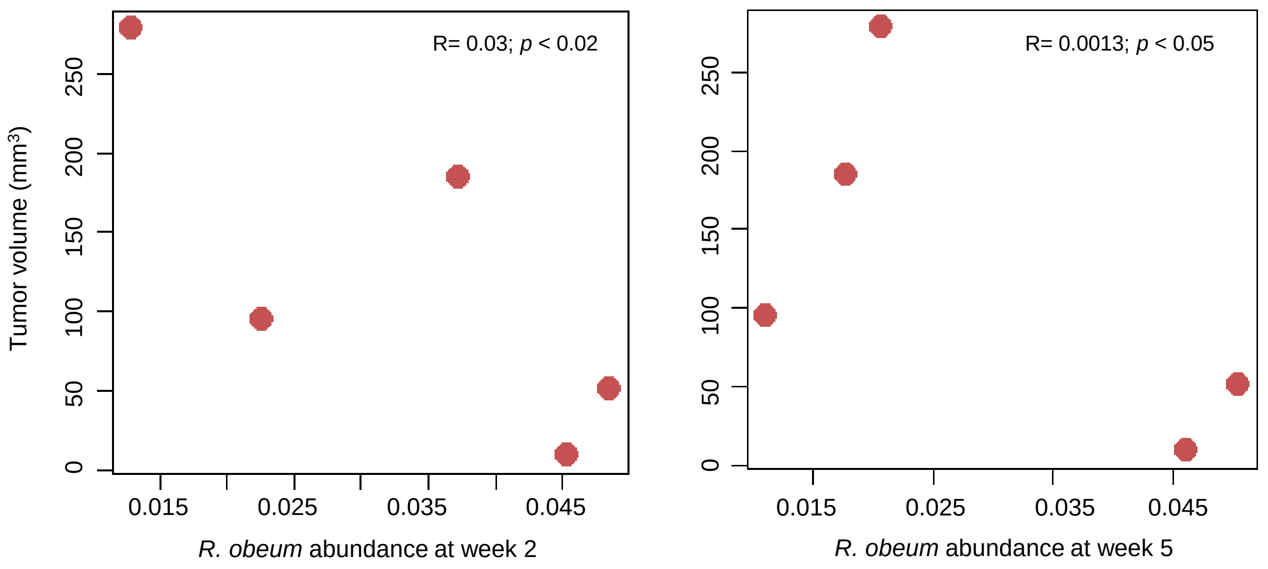


**Fig. S11**. *Ruminococcus obeum* abundance in Pathogen CAG mouse fecal samples at week 2 (left) and week 5 (right) and association with tumor volume at week 5 (only mouse fecal samples belonging to Pathogen CAG group are shown). The association between the relative abundance of *R. obeum* and tumor volume at week 5 were determined by a linear regression model after adjusting for donor (lm(= Tumor Volume at week 5 ~ Donor + *R. obeum* abundance). R= -0.69; $p < 0.036$ both time-points combined.

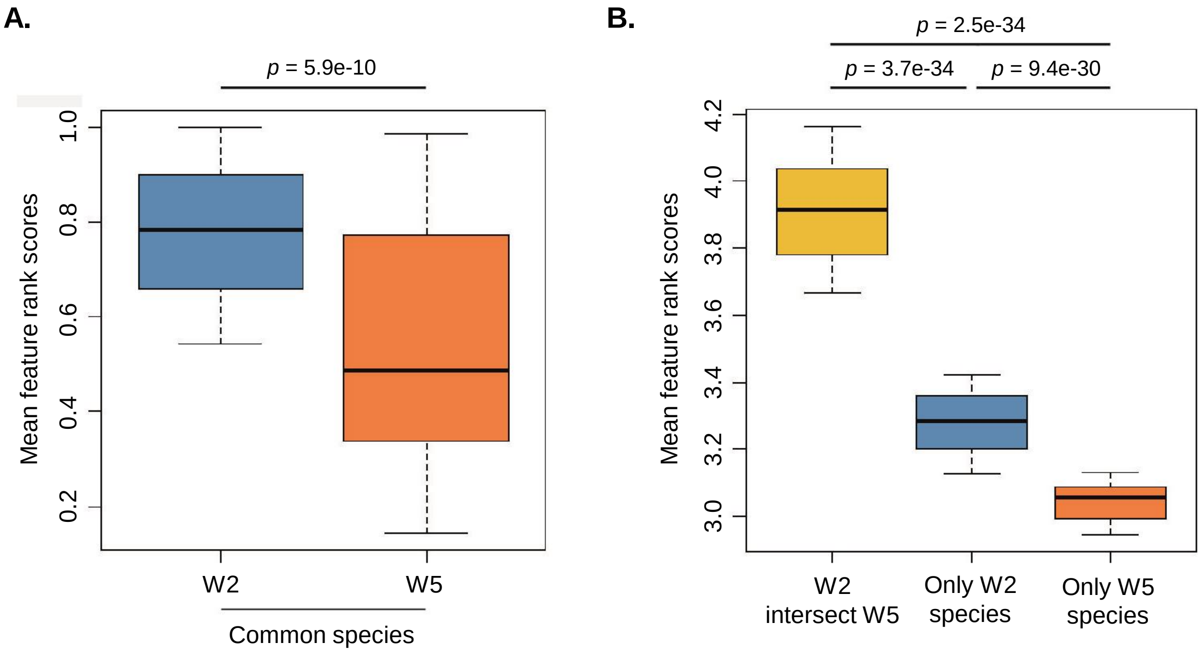


**Fig. S12. A.** Boxplot comparing the ranked feature importance scores of bacterial taxa identified as tumor-associated at both weeks 2 and 5 in the same two variants of RF models. Mann-Whitney U test *p* values for the different comparisons are indicated. **B.** Boxplots comparing the variations of the mean feature ranked scores for the three different groups (blue, abundance at week 2; orange, abundance at week 5; yellow, abundance of common taxa) of tumor-associated taxa in the disease-predictive RF models designed on the Global Reference CRC cohorts. Mann-Whitney U test *p* values for the different pairwise comparisons are indicated.

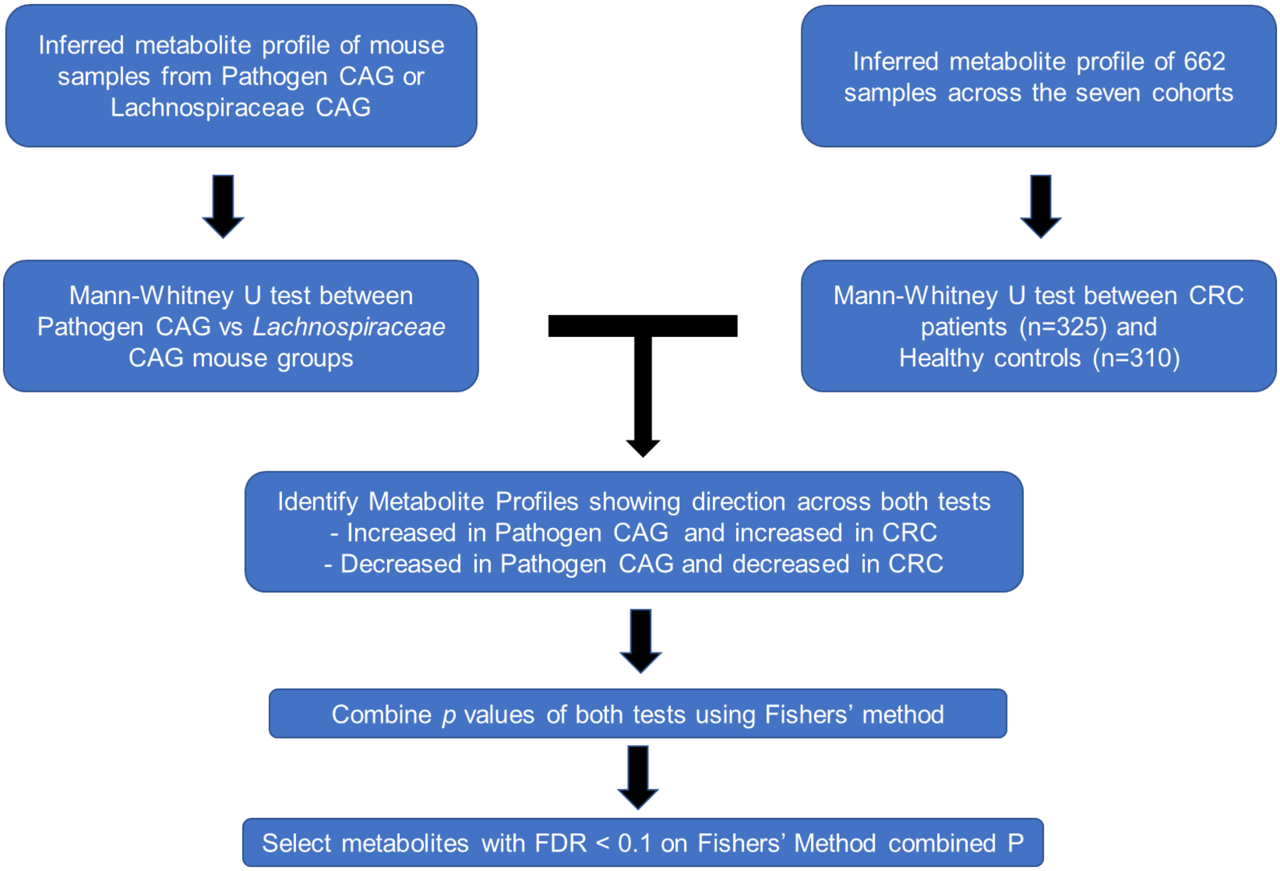


**Fig. S13.** Description of the methodology adopted for identifying a validated set of metabolic profiles that show similar patterns of association between the high-risk Pathogen and low-risk *Lachnospiraceae* microbiome types in the current study and between the CRC and healthy individuals in the Global reference CRC cohort.

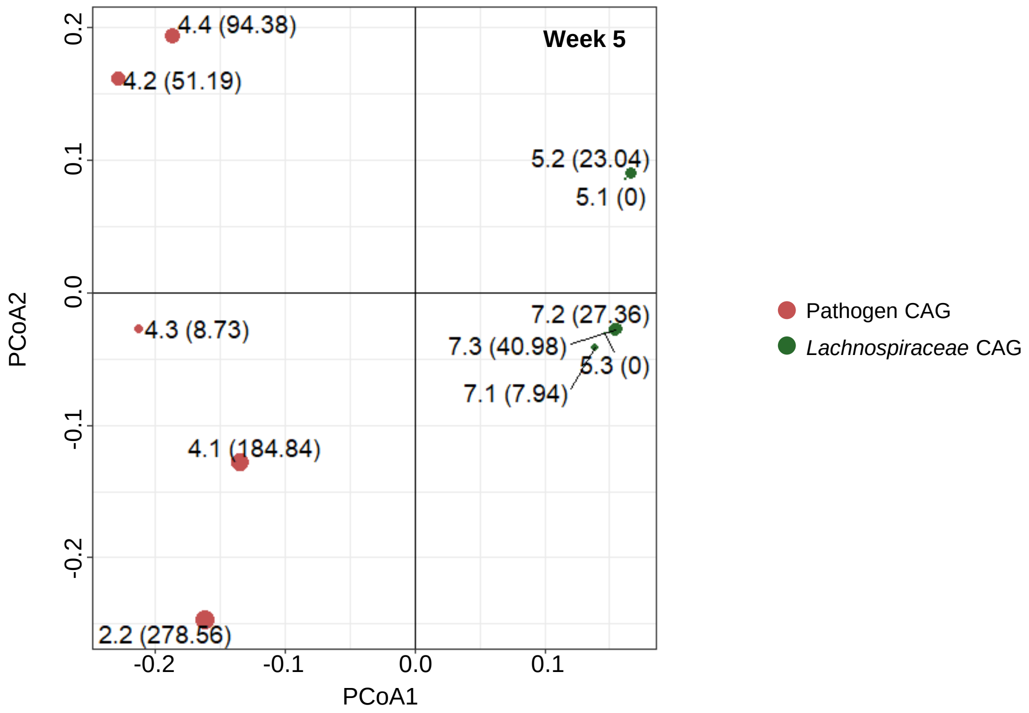


**Fig.S14.** Inferred metabolic pathway abundances predicted by AGORA/Sung Mappings differ between Pathogen CAG and *Lachnospiraceae* CAG. Principal component analysis (PCoA) based on log10 of pathway abundances. Each dot represents an individual mouse microbiota which is shown by mouse ID (tumor volume). Significant differences between the two CAG groups were calculated by PERMANOVA tests ($R^2$= 0.13; $p$ = 0.038).

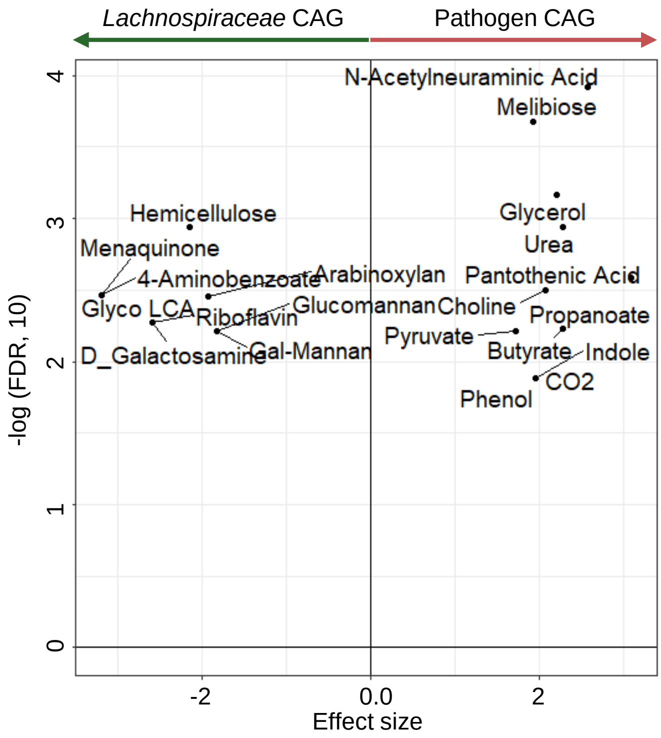


**Fig. S15.** Volcano plot showing the validated set (identified as summarized in Fig. S13) of metabolite consumption functionalities that were predicted to have either a significant positive or negative association with the Pathogen CAG microbiome. The x-axis indicates the effect size difference (negative indicating depleted in the Pathogen CAG and positive indicating enriched in the Pathogen CAG), and the y-axis indicates the negative log of FDR value.

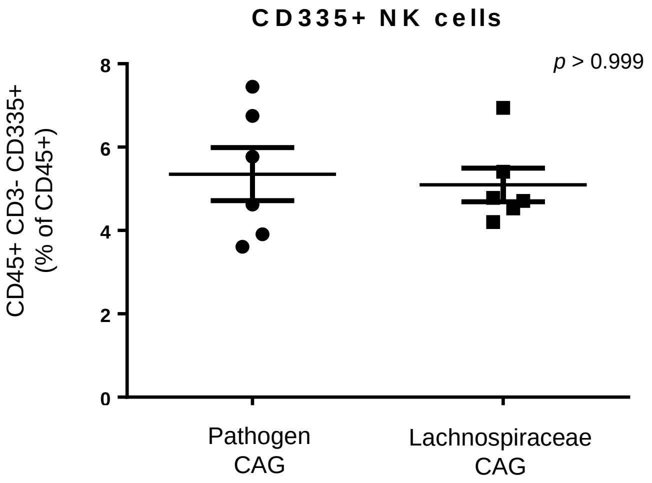


**Fig. S16.** Numbers of CD335+ NK cells are not different between mice receiving the Pathogen CAG versus *Lachnospiraceae* CAG. *p* values were determined by Mann-Whitney U test and are represented in each plot. Data indicate mean ±SEM. n = 6 biological replicates/group.

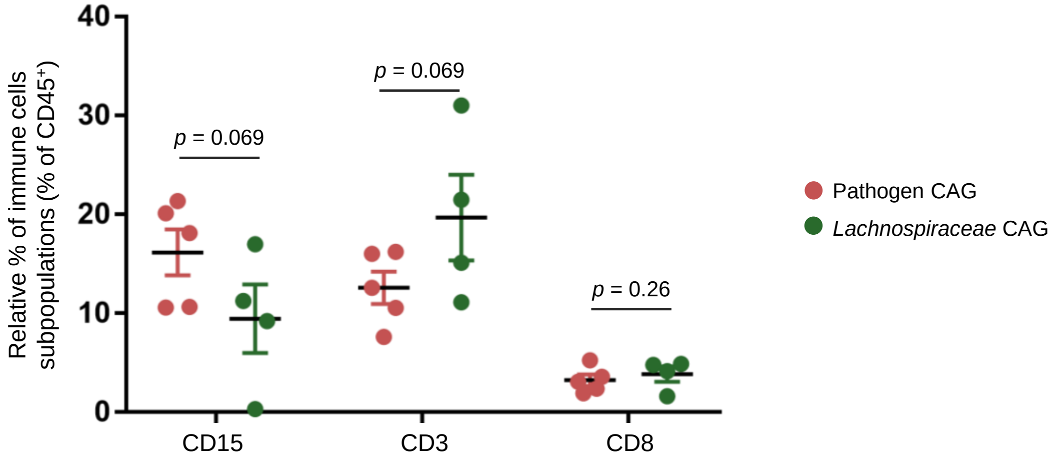


**Fig. S17.** Quantification of T-cell infiltrate ($CD3^+$ and $CD8^+$) and neutrophil infiltrate ($CD15^+$) in human CRC biopsies from Pathogen (red, n= 5) and *Lachnospiraceae*-enriched tumors (green, n= 4). 2 sections/tumor and 3 ROIs quantified per section. *p* values were calculated using unpaired t-tests between each cell type.

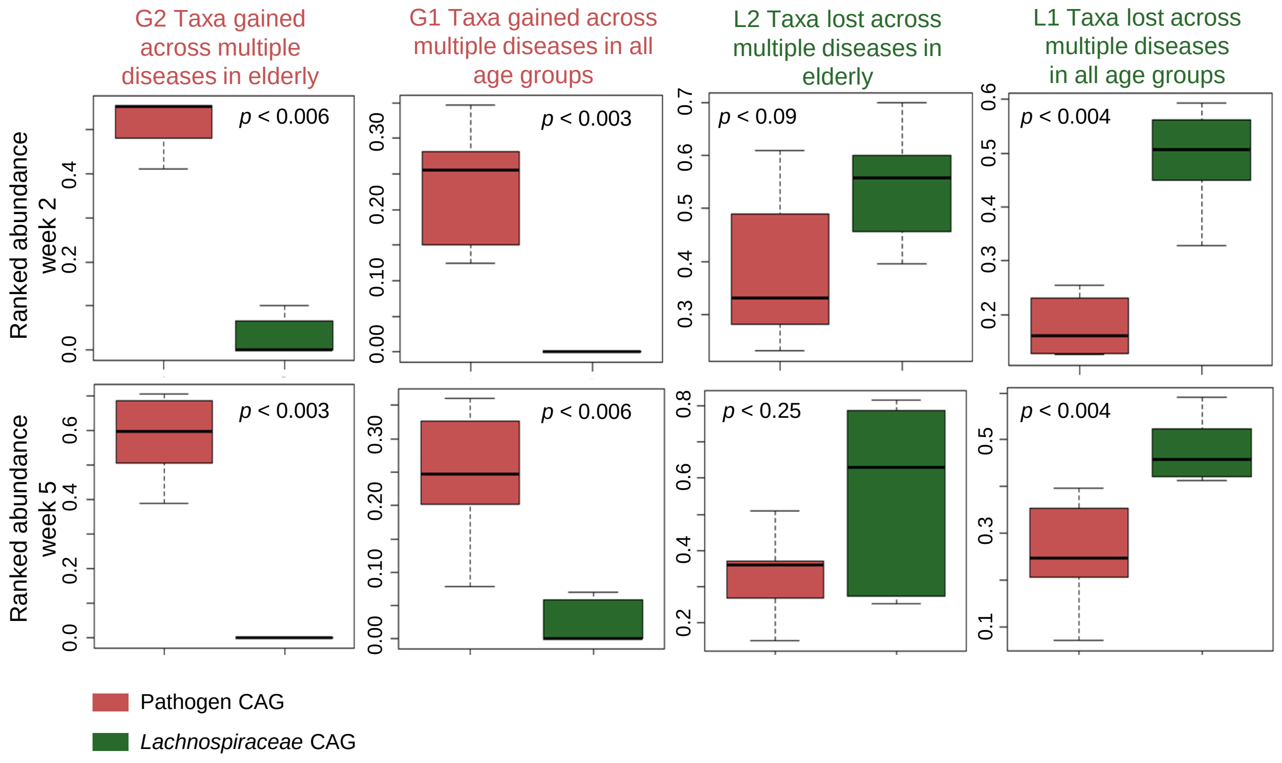


**Fig. S18.** Boxplot showing the differences in mean abundances of G2 (taxa gained across multiple diseases in elderly), G1 (taxa gained across multiple diseases across all age-groups), L2 (taxa lost across multiple diseases in elderly), and L1 (taxa lost across multiple diseases across all age-groups) taxa groups in fecal microbiomes from high-risk Pathogen CAG and low-risk *Lachnospiraceae* CAG at weeks (W) 2 and 5. Mann-Whitney U test *p* values for the different pairwise comparisons are indicated.

**Supplementary Tables**

**Table S1.** Detailed clinical and pathological information of patients with adenomas or CRC. ND, not determined; Stars (*) indicate the 12 patients selected for RNAseq analysis.

**Table S2.** CAGs abundance of 32 patients with adenomas or CRC. ON, tumor samples; UD, undiseased (normal) tissue.

**Table S3.** RNAseq statistics.

**Table S4.** Hallmark pathway analysis and CIBERSORT

**Table S5.** Human confounder analysis of four putative confounding factors (tumor type, tumor location, T category, and N category) in the RNAseq analysis. The analysis was restricted to 20 genes that were differentially expressed in Pathogen or *Lachnospiraceae* enriched tumors.

**Table S6**. Richness (Chao1, Observed species) and diversity (Shannon, Phylogenetic Diversity) of the donor was significantly higher in the *Lachnospiraceae* CAG group than in the Pathogen CAG group. *P* values were calculated with Mann-Whitney U test. The values are given as the means ± SD. The operational taxonomic units (OTUs) were defined with 97% sequence identity.  Calculations were made based on rarefied OTU table at 11386 sequences.

**Table S7**. Details of the shallow shotgun data analysis.

**Table S8**. Description of the predicted metabolites profile and the corresponding bacterial species.

| Genes | Tumor Type | | Tumor Location | | T-Category | | N-Category | |
|---|---|---|---|---|---|---|---|---|
| | F-value | FDR | F-value | FDR | F-value | FDR | F-value | FDR |
| CSF1 | **10.63** | **0.08** | 3.80 | 1.00 | **14.01** | **0.04** | 0.63 | 1.00 |
| IL1A | 3.07 | 1.00 | 1.85 | 1.00 | 2.68 | 1.00 | 0.06 | 1.00 |
| CTLA4 | 5.11 | 0.59 | 0.29 | 1.00 | 1.25 | 1.00 | 0.46 | 1.00 |
| CXCR2 | 1.18 | 1.00 | 0.09 | 1.00 | 0.52 | 1.00 | 1.04 | 1.00 |
| PDCD1 | 0.81 | 1.00 | 0.28 | 1.00 | 0.39 | 1.00 | 0.18 | 1.00 |
| PSMD2 | 2.11 | 1.00 | 3.79 | 1.00 | 3.90 | 1.00 | 0.19 | 1.00 |
| CXCL11 | 0.59 | 1.00 | 1.28 | 1.00 | 1.09 | 1.00 | 1.27 | 1.00 |
| CXCL13 | 1.11 | 1.00 | 0.74 | 1.00 | 0.48 | 1.00 | 0.22 | 1.00 |
| CSF1R | 3.67 | 1.00 | 1.55 | 1.00 | 2.19 | 1.00 | 0.49 | 1.00 |
| HLA-C | 0.21 | 1.00 | 0.40 | 1.00 | 2.93 | 1.00 | 0.13 | 1.00 |
| HLA-B | 2.07 | 1.00 | 2.19 | 1.00 | 1.13 | 1.00 | 0.09 | 1.00 |
| ULBP3 | 1.62 | 1.00 | 3.27 | 1.00 | 1.94 | 1.00 | 0.34 | 1.00 |
| IL33 | **12.29** | **0.05** | **9.68** | **0.10** | 4.69 | 0.71 | 1.01 | 1.00 |
| NLRP6 | 0.89 | 1.00 | 0.45 | 1.00 | 0.04 | 1.00 | 0.79 | 1.00 |
| CD3E | 0.94 | 1.00 | 0.66 | 1.00 | 0.81 | 1.00 | 0.33 | 1.00 |
| MAP3K8 | 0.09 | 1.00 | 0.20 | 1.00 | 0.20 | 1.00 | 0.39 | 1.00 |
| CD163 | 3.69 | 1.00 | 1.41 | 1.00 | 2.98 | 1.00 | 1.43 | 1.00 |
| MAP3K14 | 0.79 | 1.00 | 0.82 | 1.00 | 0.85 | 1.00 | 1.03 | 1.00 |
| TRPC4AP | 1.07 | 1.00 | 9.15 | 0.11 | 0.81 | 1.00 | 3.34 | 1.00 |
| CBLC | 2.24 | 1.00 | 1.77 | 1.00 | 1.24 | 1.00 | 1.40 | 1.00 |

**Table S5.** Human confounder analysis of four putative confounding factors (tumor type, tumor location, T category, and N category) in the RNAseq analysis. The analysis was restricted to 20 genes that were differentially expressed in Pathogen or *Lachnospiraceae* enriched tumors.

| Alpha diversity measures | Pathogen CAG | Lachnospiraceae CAG | *p* value* |
|---|---|---|---|
| **Week 1** | | | |
| Chao1 | 141.2 ± 14.1 | 184.2 ± 27.6 | 0.009 |
| Phylogenetic diversity | 8.6 ± 0.7 | 12.3 ± 1.4 | 0.002 |
| Shannon index | 3.5 ± 0.3 | 4.7 ± 0.2 | 0.002 |
| Observed species | 109.5 ± 6.9 | 146.3 ± 10.1 | 0.002 |
| **Week 2** | | | |
| Chao1 | 187.3 ± 38.3 | 276.6 ± 8.8 | 0.002 |
| Phylogenetic diversity | 11.1 ± 1.5 | 16.7 ± 0.6 | 0.002 |
| Shannon index | 4.2 ± 0.1 | 4.8 ± 0.2 | 0.002 |
| Observed species | 137.8 ± 11.5 | 202.8 ± 12.6 | 0.005 |
| **Week 3** | | | |
| Chao1 | 181.6 ± 31.2 | 257.3 ± 26.4 | 0.004 |
| Phylogenetic diversity | 11.3 ± 1.2 | 16.4 ± 0.7 | 0.002 |
| Shannon index | 4.3 ± 0.2 | 4.7 ± 0.2 | 0.026 |
| Observed species | 144 ± 10.1 | 197.3 ± 9 | 0.005 |
| **Week 4** | | | |
| Chao1 | 192.7 ± 18.1 | 254.7 ± 30.1 | 0.002 |
| Phylogenetic diversity | 11.5 ± 1 | 16.5 ± 1.1 | 0.002 |
| Shannon index | 4.4 ± 0.3 | 5 ± 0.1 | 0.009 |
| Observed species | 149.2 ± 10.4 | 207.5 ± 12.9 | 0.002 |
| **Week 5** | | | |
| Chao1 | 217.4 ± 27.3 | 247.9 ± 19.8 | 0.026 |
| Phylogenetic diversity | 12 ± 1.3 | 16.2 ± 1.2 | 0.002 |
| Shannon index | 4.4 ± 0.4 | 5.1 ± 0.1 | 0.002 |
| Observed species | 159.3 ± 13.7 | 203.8 ± 8.6 | 0.002 |

**Table S6**. Richness (Chao1, Observed species) and diversity (Shannon, Phylogenetic Diversity) of the donor was significantly higher in the *Lachnospiraceae* CAG group than in the Pathogen CAG group. *P* values were calculated with Mann-Whitney U test. The values are given as the means ± SD. The operational taxonomic units (OTUs) were defined with 97% sequence identity.  Calculations were made based on rarefied OTU table at 11386 sequences.

### Metabolites Consumption

| | Trimethylamine | p_Cresol | NH3 | CO2 | Acetone | Phenylacetate | Indole | Lithocholic | Deoxycholic | Folic | Chenodeoxycholic | Adenosylcobalamin | Thiamine | Cholic | Pyridoxal | Cohen's D (Donor1 vs Donor2) |
|---|---|---|---|---|---|---|---|---|---|---|---|---|---|---|---|---|
| Clostridium_symbiosum | 0 | 1 | 1 | 1 | 1 | 0 | 0 | 0 | 0 | 0 | 0 | 0 | 0 | 0 | 0 | 2.74 |
| Parabacteroides_distasonis | 0 | 0 | 0 | 0 | 0 | 1 | 0 | 0 | 0 | 0 | 0 | 0 | 0 | 0 | 0 | 2.43 |
| Clostridium_hathewayi | 1 | 1 | 1 | 1 | 1 | 0 | 0 | 0 | 0 | 0 | 0 | 0 | 0 | 0 | 0 | 2.06 |
| Bacteroides_caccae | 0 | 1 | 0 | 0 | 0 | 0 | 0 | 0 | 0 | 0 | 0 | 0 | 0 | 0 | 0 | 2.04 |
| Bacteroides_salyersiae | 0 | 0 | 0 | 0 | 0 | 0 | 1 | 0 | 0 | 0 | 0 | 0 | 0 | 0 | 0 | 1.97 |
| Bacteroides_fragilis | 0 | 0 | 0 | 0 | 0 | 1 | 1 | 1 | 1 | 0 | 0 | 0 | 0 | 0 | 0 | 1.93 |
| Bacteroides_uniformis | 0 | 1 | 0 | 0 | 0 | 1 | 1 | 0 | 0 | 0 | 0 | 0 | 0 | 0 | 0 | 1.71 |
| Clostridium_bolteae | 0 | 0 | 1 | 1 | 1 | 0 | 0 | 0 | 0 | 0 | 0 | 0 | 0 | 0 | 0 | 1.35 |
| Clostridium_hylemonae | 0 | 0 | 1 | 1 | 1 | 0 | 0 | 1 | 1 | 0 | 0 | 0 | 0 | 0 | 0 | 1.07 |
| Clostridium_asparagiforme | 1 | 1 | 1 | 1 | 1 | 0 | 0 | 0 | 0 | 0 | 0 | 0 | 0 | 0 | 0 | 0.83 |
| Bifidobacterium_breve | 0 | 0 | 0 | 0 | 0 | 0 | 0 | 1 | 1 | 1 | 1 | 1 | 1 | 1 | 1 | 0.66 |
| Alistipes_putredinis | 0 | 0 | 0 | 0 | 0 | 0 | 1 | 0 | 0 | 0 | 0 | 0 | 0 | 0 | 0 | -1.18 |
| Odoribacter_splanchnicus | 0 | 0 | 0 | 0 | 0 | 0 | 1 | 0 | 0 | 0 | 0 | 0 | 0 | 0 | 0 | -1.90 |
| Bifidobacterium_pseudocatenulatum | 0 | 0 | 0 | 0 | 0 | 0 | 0 | 0 | 0 | 1 | 0 | 0 | 0 | 0 | 0 | -2.30 |
| Bacteroides_thetaiotaomicron | 0 | 0 | 0 | 0 | 0 | 1 | 1 | 0 | 0 | 0 | 0 | 0 | 0 | 0 | 0 | -2.87 |
| Bifidobacterium_longum | 0 | 0 | 0 | 0 | 0 | 0 | 0 | 0 | 0 | 1 | 1 | 1 | 1 | 1 | 1 | -3.19 |
| Bacteroides_dorei | 0 | 1 | 0 | 0 | 0 | 0 | 0 | 0 | 0 | 0 | 0 | 0 | 0 | 0 | 0 | -3.35 |
| Bilophila_wadsworthia | 0 | 0 | 1 | 0 | 0 | 0 | 0 | 0 | 1 | 0 | 0 | 0 | 0 | 0 | 0 | -3.63 |
| Alistipes_onderdonkii | 0 | 0 | 0 | 0 | 0 | 0 | 1 | 0 | 0 | 0 | 0 | 0 | 0 | 0 | 0 | -3.77 |

### Metabolites Production

| | Pantothenic Acid | Urea | Arabinoxylan | Hemicellulose | N_Acetylneuraminic Acid | Melibiose | Pyruvate | Glycerol | Menaquinone | X4_Aminobenzoate | D_Galactosamine | Glycolithocholate | Galactomannan | Glucomannan | Choline | Butyrate | Indole | Propanoate | Phenol | CO2 | Cohen's D (Donor1 vs Donor2) |
|---|---|---|---|---|---|---|---|---|---|---|---|---|---|---|---|---|---|---|---|---|---|
| Clostridium_symbiosum | 1 | 1 | 0 | 0 | 1 | 0 | 1 | 0 | 0 | 0 | 0 | 0 | 0 | 0 | 0 | 0 | 0 | 0 | 0 | 0 | 2.74 |
| Parabacteroides_distasonis | 0 | 0 | 0 | 1 | 0 | 1 | 0 | 0 | 0 | 0 | 0 | 0 | 0 | 0 | 0 | 0 | 0 | 0 | 0 | 0 | 2.43 |
| Desulfovibrio_piger | 0 | 1 | 0 | 0 | 0 | 0 | 1 | 1 | 0 | 0 | 0 | 0 | 0 | 0 | 0 | 1 | 1 | 1 | 1 | 1 | 2.28 |
| Clostridium_hathewayi | 1 | 1 | 0 | 0 | 0 | 0 | 0 | 0 | 0 | 0 | 0 | 0 | 0 | 0 | 1 | 0 | 0 | 0 | 0 | 0 | 2.06 |
| Bacteroides_caccae | 0 | 0 | 0 | 0 | 1 | 1 | 0 | 0 | 0 | 0 | 0 | 0 | 0 | 0 | 0 | 0 | 0 | 0 | 0 | 0 | 2.04 |
| Bacteroides_fragilis | 0 | 0 | 0 | 1 | 1 | 1 | 0 | 0 | 0 | 0 | 0 | 0 | 0 | 0 | 0 | 0 | 0 | 0 | 0 | 0 | 1.93 |
| Bacteroides_xylanisolvens | 0 | 0 | 0 | 0 | 0 | 0 | 0 | 1 | 0 | 0 | 0 | 0 | 0 | 0 | 0 | 0 | 0 | 0 | 0 | 0 | 1.86 |
| Bacteroides_uniformis | 0 | 0 | 0 | 0 | 0 | 1 | 0 | 0 | 0 | 0 | 0 | 0 | 1 | 1 | 0 | 0 | 0 | 0 | 0 | 0 | 1.71 |
| Clostridium_bolteae | 1 | 1 | 0 | 0 | 0 | 0 | 0 | 1 | 0 | 0 | 0 | 0 | 0 | 0 | 0 | 0 | 0 | 0 | 0 | 0 | 1.35 |
| Clostridium_hylemonae | 1 | 1 | 0 | 0 | 0 | 0 | 0 | 0 | 0 | 0 | 0 | 0 | 0 | 0 | 0 | 0 | 0 | 0 | 0 | 0 | 1.07 |
| Clostridium_asparagiforme | 1 | 1 | 0 | 0 | 0 | 0 | 0 | 0 | 0 | 0 | 0 | 0 | 0 | 0 | 1 | 0 | 0 | 0 | 0 | 0 | 0.83 |
| Bifidobacterium_breve | 0 | 1 | 0 | 0 | 1 | 1 | 0 | 0 | 0 | 0 | 1 | 1 | 0 | 0 | 0 | 0 | 0 | 0 | 0 | 0 | 0.66 |
| Roseburia_inulinivorans | 0 | 0 | 0 | 1 | 0 | 1 | 0 | 0 | 0 | 0 | 0 | 0 | 0 | 0 | 0 | 0 | 0 | 0 | 0 | 0 | -0.78 |
| Collinsella_aerofaciens | 0 | 0 | 0 | 0 | 1 | 0 | 0 | 0 | 0 | 0 | 0 | 0 | 0 | 0 | 0 | 0 | 0 | 0 | 0 | 0 | -1.82 |
| Akkermansia_muciniphila | 0 | 0 | 0 | 0 | 1 | 0 | 0 | 0 | 0 | 0 | 0 | 0 | 0 | 0 | 0 | 0 | 0 | 0 | 0 | 0 | -1.88 |
| Bifidobacterium_pseudocatenulatum | 0 | 1 | 1 | 1 | 0 | 0 | 0 | 0 | 0 | 0 | 0 | 0 | 0 | 0 | 0 | 0 | 0 | 0 | 0 | 0 | -2.30 |
| Bacteroides_thetaiotaomicron | 0 | 0 | 1 | 1 | 0 | 1 | 0 | 0 | 0 | 0 | 0 | 0 | 0 | 0 | 0 | 0 | 0 | 0 | 0 | 0 | -2.87 |
| Bifidobacterium_longum | 0 | 1 | 1 | 1 | 1 | 1 | 0 | 1 | 1 | 1 | 1 | 1 | 0 | 0 | 0 | 0 | 0 | 0 | 0 | 0 | -3.19 |
| Bacteroides_dorei | 0 | 0 | 0 | 1 | 0 | 0 | 0 | 0 | 0 | 0 | 0 | 0 | 0 | 0 | 0 | 0 | 0 | 0 | 0 | 0 | -3.35 |
| Bilophila_wadsworthia | 0 | 0 | 0 | 0 | 0 | 0 | 1 | 1 | 0 | 0 | 0 | 0 | 0 | 0 | 0 | 0 | 0 | 0 | 0 | 0 | -3.63 |
| Bacteroides_cellulosilyticus | 0 | 0 | 1 | 1 | 0 | 0 | 0 | 0 | 0 | 0 | 0 | 0 | 1 | 1 | 0 | 0 | 0 | 0 | 0 | 0 | -8.46 |

**Table S8**. Description of the predicted metabolites profile and the corresponding bacterial species.
